# Drug addiction mutations unveil a methylation ceiling in EZH2-mutant lymphoma

**DOI:** 10.1101/2022.04.04.486977

**Authors:** Hui Si Kwok, Allyson M. Freedy, Allison P. Siegenfeld, Julia W. Morriss, Amanda L. Waterbury, Stephen M. Kissler, Brian B. Liau

**Affiliations:** Department of Chemistry and Chemical Biology, Harvard University, Cambridge, MA 02138, USA; Broad Institute of MIT and Harvard, Cambridge, MA 02142, USA; Department of Immunology and Infectious Diseases, Harvard T.H. Chan School of Public Health, Boston, MA 02115, USA

## Abstract

Cancer mutations in Polycomb Repressive Complex 2 (PRC2) drive aberrant epigenetic states. Although therapies inhibiting the PRC2 enzymatic component EZH2 are FDA-approved, oncogene-specific dependencies remain to be discovered. Here, we identify mutations that confer both resistance and drug addiction to PRC2 inhibitors in EZH2-mutant lymphoma, resulting in cancer cells that paradoxically depend on drug for survival. Drug addiction is mediated by hypermorphic mutations in the CXC domain of EZH2, which maintain H3K27me3 levels even in the presence of PRC2 inhibitors. Drug removal leads to overspreading of H3K27me3, surpassing a repressive methylation ceiling compatible with lymphoma cell survival. Activating EZH2 cancer mutations establish an epigenetic state precariously close to this ceiling, which we show can be breached by inhibition of SETD2, a PRC2 antagonist, to block lymphoma growth. More broadly, we highlight how approaches to identify drug addiction mutations can be leveraged to discover cancer vulnerabilities.

## Main Text

Recurrent activating mutations in EZH2, the catalytic subunit of the transcriptional repressor PRC2, can promote neoplastic growth by silencing tumor suppressor genes in a wide range of lymphomas, including roughly 20% of follicular lymphomas (FLs) and 10% of diffuse large B-cell lymphomas (DLBCLs) (*1*). The discovery of these activating mutations prompted the development of PRC2 small molecule inhibitors, leading to the approval of the EZH2 active-site inhibitor tazemetostat for FL and the clinical development of allosteric inhibitors targeting the PRC2 subunit, EED (*2*). Both active and allosteric site PRC2 inhibitors reduce H3K27me3 levels and derepress tumor suppressor genes (*3*–*5*). Cellular models confirmed that lymphomas can develop resistance to PRC2 inhibitors by acquiring mutations in EZH2 that disrupt drug binding and restore H3K27me3 levels (*6, 7*). How resistance might emerge to these therapies in the clinic has yet to be fully determined. Consequently, a comprehensive map of the genetic and biochemical basis of resistance to PRC2 inhibitors could improve the clinical management of lymphomas and enhance our basic understanding of PCR2 regulation and EZH2-mutant co-dependencies (*8*). In particular, such studies could reveal if the increased levels of H3K27me3 in EZH2-mutant lymphomas may present distinct vulnerabilities beyond PRC2 inhibition.

To investigate these possibilities, we surveyed the drug resistance landscape to PRC2 inhibitors in EZH2-mutant lymphoma using CRISPR-suppressor scanning, a method we previously developed to identify drug resistance hot spots within a protein target (*9*). We performed CRISPR-suppressor scanning in the DLBCL cell line Karpas-422, which is heterozygous for the most common activating EZH2 mutation, *EZH2* Y646N (*10*). We used a pool of 650 sgRNAs spanning the coding sequence of the core members of PRC2 (EZH2, EED and SUZ12) and profiled resistance to two PRC2 inhibitors: GSK343, an EZH2 active site inhibitor (*11*); and EED226, an EED allosteric inhibitor that blocks PRC2 allosteric activation by H3K27me3 (**Fig. 1A, Supplementary Data S1**) (*4, 5*). Following 5 weeks of inhibitor treatment, we isolated genomic DNA from surviving cells and determined the enrichment of sgRNAs under each treatment condition. The top-enriched sgRNAs unique to each drug treatment (EZH2 sgI109, EZH2 sgC699, and EED sgY365) targeted positions within the drug binding sites of the respective inhibitors (**Figs. 1B-D, S1, S2A-B, Supplementary Data S2**) (*4, 12*). Sequencing of the exons encompassing these positions revealed an enrichment of mutations that likely disrupt drug binding (*EZH2* I109_M110insM under GSK343 and *EED* Y365F/M366V under EED226 treatment, **Fig. S2C-D**) (*6, 7, 12*).

**Fig. 1.**
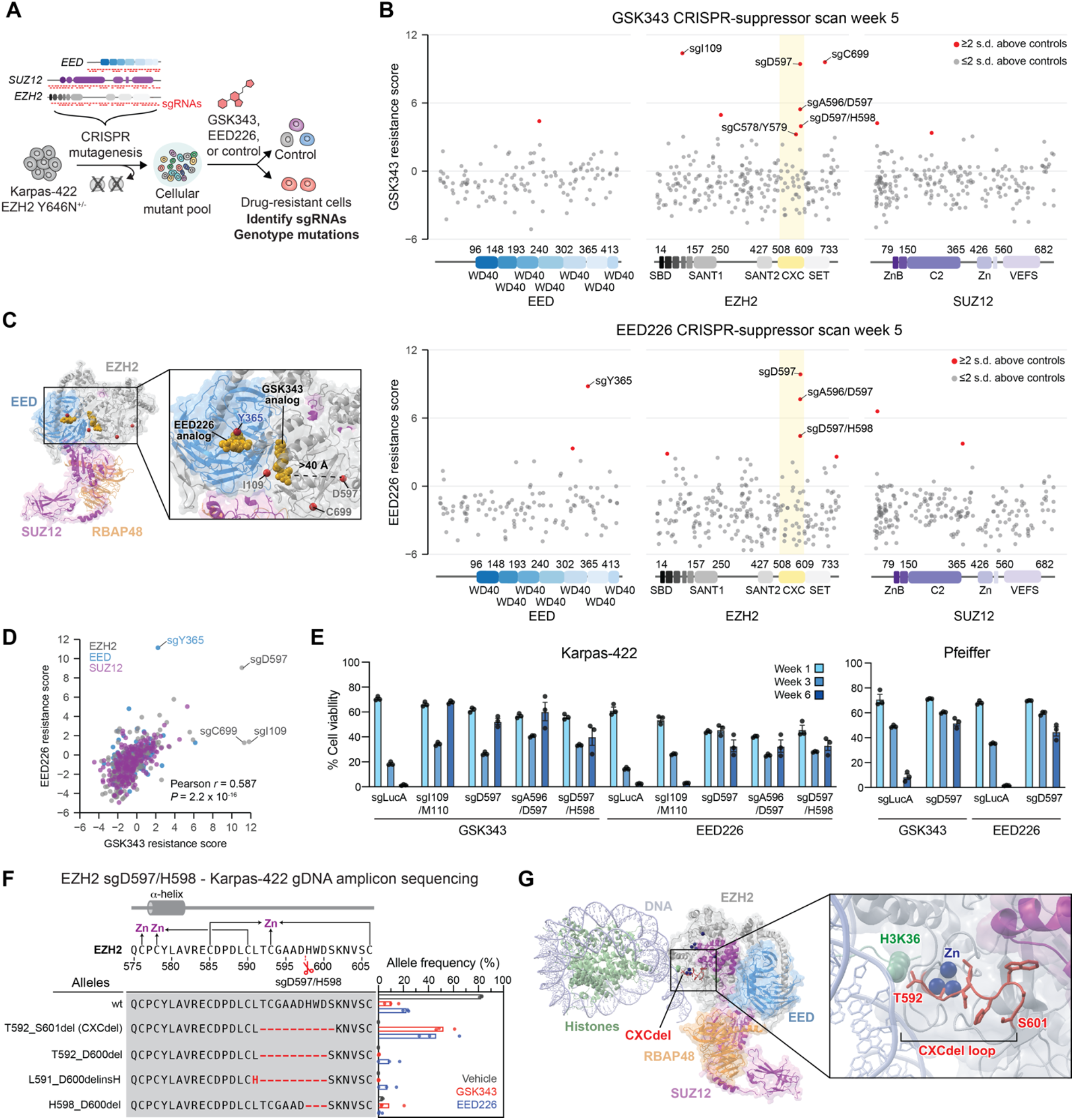
CRISPR-suppressor scanning identifies the EZH2 CXC domain as a drug resistance hotspot to PRC2 inhibitors. **A**. Schematic showing the CRISPR-suppressor scanning workflow applied to profile resistance mutations in PRC2 (EED, SUZ12 and EZH2) to inhibitors, GSK343 and EED226. Cells were transduced with the PRC2-targeting sgRNA library and treated with PRC2 inhibitors or vehicle for 5 weeks. sgRNA enrichment frequencies were determined by sequencing genomic DNA (gDNA) from the surviving cellular pool. **B**. Scatter plots showing resistance scores (*y-*axis) in Karpas-422 under 1 μM GSK343 treatment (top) and 1 μM EED226 treatment (bottom) at week 5. Resistance scores were calculated as the log_2_(fold-change) sgRNA enrichment under drug normalized to the mean of the negative control sgRNAs (*n* = 58). The PRC2-targeting guides (*n* = 650) are arrayed by amino acid position in the EED, EZH2 or SUZ12 coding sequence on the *x*-axis corresponding to the position of the predicted cut site. When the sgRNA cut site falls between two amino acids, both amino acids are denoted. Data points represent mean value across three replicates. Protein domains are demarcated by colored panels along the *x*-axis. The location of the EZH2 CXC domain is shown by a yellow background. Points colored in red have resistance scores greater than 2 standard deviations (s.d.) above the mean of the negative control sgRNAs. **C**. Structural view showing enriched sgRNAs in EZH2 and EED drug binding sites identified in the GSK343 and EED226 CRISPR-suppressor scanning experiments. GSK343 and EED226 are shown in gold. Amino acids corresponding to enriched sgRNAs are shown as red spheres. PDB: 6WKR, 5K0M, 5LS6 **D**. Scatter plot showing sgRNA resistance scores in the EED226 CRISPR-suppressor scan (*y-* axis) and GSK343 CRISPR-suppressor scan (*x-*axis). **E**. Bar plots of cell viability over a time course of 6 weeks following lentiviral transduction of individual sgRNAs and treatment with GSK343 (1 μM for Karpas-422 and incremental doses up to 200 nM for Pfeiffer) or EED226 (1 μM for Karpas-422 and incremental doses up to 500 nM for Pfeiffer) in Karpas-422 (left) and Pfeiffer cells (right). Bars represent mean ± s.e.m across three replicates. **F**. Schematic showing genotypes and bar plots of allele frequencies for mutations that are observed at frequencies of >5% in the gDNA encoding *EZH2* surrounding the cut site of CXC-targeting sgRNAs for GSK343 (red) or EED226 (blue) treatment at week 6 in Karpas-422 cells transduced with sgD597/H598. (top) Schematic depicts the secondary structure of C-terminal CXC domain and cysteine residues that coordinate the zinc ions. **G**. Structural view of PRC2 engaged with a mononucleosome showing the location of CXCdel loop (red) and zinc ions (blue). EMDB-7306, PDB: 6WKR

Unexpectedly, several sgRNAs targeting positions within the CXC domain of EZH2 were also highly enriched in both drug treatment conditions (sgA596/D597, sgD597, sgD597/H598, **Figs. 1B-D, S1, Supplementary Data S2**). These sgRNAs target a loop in the CXC domain that lies 17 Å from the substrate nucleosome and over 40 Å from either inhibitor binding site (**Fig. 1C**) (*4, 12*–*14*). When individually transduced in Karpas-422 cells, these top-enriched EZH2 CXC domain sgRNAs conferred robust resistance to both drug treatments, while cells transduced with EZH2 sgI109/M110 only survived in the presence of GSK343 (**Fig. 1E**), consistent with our CRISPR-suppressor scan data. Additionally, transduction of EZH2 sgD597 also conferred robust resistance to PRC2 inhibitors in Pfeiffer, a DLBCL cell line heterozygous for the PRC2 activating mutation *EZH2* A682G (**Fig. 1E**). Taken together, these results demonstrate that mutations in the CXC domain of EZH2 confer drug resistance through an unknown mechanism in multiple DLBCL cell lines with distinct EZH2-activating mutations.

Genotyping the resultant bulk drug resistant cells from individual CXC-targeting sgRNA transductions revealed frequent in-frame deletion mutations within the EZH2 CXC domain (**Figs. 1F-G, S3**). The most highly enriched mutation was a 10-amino acid deletion spanning EZH2 T592–S601, which was identified in all three CXC-targeting sgRNAs transductions across both drug treatments. The EZH2 T592–S601 deletion modifies a poorly conserved loop adjacent to a cluster of cysteine residues that coordinate zinc ions (**Figs. 1F, S4**) (*15*). Hereafter, this mutation is referred to as CXCdel. Notably, *EZH2* CXCdel is not enriched in the absence of PRC2 inhibitors (**Figs. 1G, S3**), suggesting that drug treatment strongly selects for the deletion mutation.

We sought to generate clonal Karpas-422 CXCdel cells. Despite the strong enrichment of *EZH2* CXCdel in the presence of PRC2 inhibitors, clonal cell lines containing CXCdel could not be expanded from single cells when sorted into fresh media lacking inhibitor. We considered if CXCdel mutant cells have a growth defect in the absence of drug. To test this possibility, the PRC2 CRISPR-suppressor scan was repeated and further extended by 3 weeks either in the presence of PRC2 inhibitors or after their withdrawal (**Fig. 2A**). Through this adapted method, which we term ‘CRISPR-addiction scanning’, we observed that the previously enriched CXC-targeting sgRNAs were highly depleted upon drug removal, while the sgRNAs targeting the respective drug binding sites remained enriched (**Figs. 2B, S5**). Accordingly, genotyping revealed that *EZH2* CXCdel was depleted upon drug withdrawal (**Fig. S6**), consistent with the hypothesis that cells with this mutation are drug addicted (*i*.*e*., have reduced intrinsic fitness upon drug removal) (*16, 17*).

**Fig. 2.**
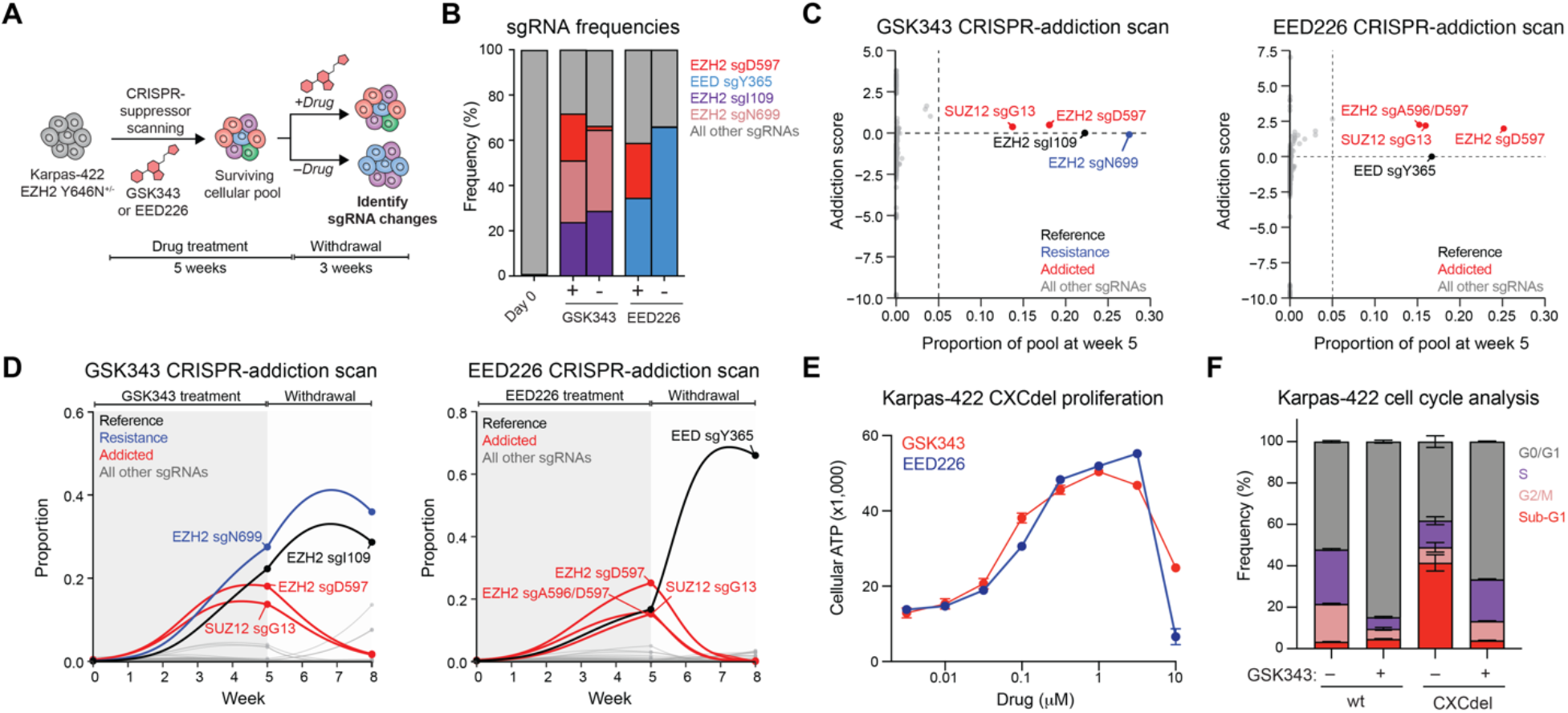
CRISPR-addiction scanning reveals that mutations in the EZH2 CXC domain confer addiction to PRC2 inhibitors. **A**. Schematic showing the CRISPR-addiction scanning workflow applied to identify drug addiction mutations in PRC2 (EED, SUZ12 and EZH2) to GSK343 and EED226. Cells transduced with the sgRNA library were selected with PRC2 inhibitors for 5 weeks and then split into two parallel treatment conditions: (1) under 3 additional weeks of continuous drug treatment and (2) 3 weeks of drug withdrawal. sgRNA enrichment frequencies were then determined by sequencing gDNA from the surviving cellular pool. **B**. Stacked bar chart showing representation of top-enriched sgRNAs (>20% in frequency) in the surviving mutant pool under continuous GSK343 or EED226 (1 μM) treatments for 8 weeks versus 5 weeks of continuous drug treatment followed by 3 weeks of drug removal. **C**. Scatter plot showing addiction score (*y*-axis) versus proportion of the sgRNA pool made up by each sgRNA at week 5 (*x*-axis) for the GSK343 CRISPR-addiction scan (left) and EED226 CRISPR-addiction scan (right). Proportion of the sgRNA pool was averaged across the three replicates of the experiment and the addiction score was calculated as detailed in Supplementary Text. **D**. Graph showing proportions of each sgRNA (*y-*axis) in the GSK343 CRISPR-addiction scan (left) and EED226 CRISPR-addiction scan (right) over time (*x*-axis). The grey background indicates the treatment duration of GSK343 or EED226 and white background indicates the period of drug removal. Curves were calculated using the estimated intrinsic growth rate *r* for each sgRNA-containing cell population under the assumption of competitive logistic growth. The reference sgRNA is shown in black, sgRNAs demonstrating resistance without addiction are shown in blue, and addicted sgRNAs are shown in red. **E**. Dose-response proliferation curves of Karpas-422 CXCdel under GSK343 and EED226 treatment for 10 days. Data represent mean ± s.e.m across three replicates. **F**. Cell cycle analysis of wt and CXCdel Karpas-422 cells under 1 μM GSK343 or vehicle treatment for 7 days. Data represent mean ± s.e.m across three replicates.

An alternative hypothesis for the depletion of CXC-targeting sgRNAs is that cells containing these sgRNAs were out-competed upon drug removal by other cells containing previously suppressed sgRNAs (*i*.*e*., a reduction in the relative fitness of CXC-targeting sgRNAs, despite no change in intrinsic fitness). To test this hypothesis, we fit a system of ordinary differential equations describing competitive logistic growth (the competitive Lotka-Volterra equations; see **Supplemental Text**) to the sgRNA prevalence data at weeks 0, 5, and 8 (*18*). By comparing the estimated intrinsic growth rate *r* for each sgRNA-containing population on-drug (*r*_*on*_) *vs*. off-drug (*r*_*off*_) relative to a drug-resistant mutant that disrupts inhibitor binding with intrinsic fitness that is assumed to be unaffected by drug (*i*.*e*., EZH2 sgI109 for GSK343 and EED sgY365 for EED226), we derived an addiction score, *A* = *r*_*on*_ – *r*_*off*_. This score is positive when the sgRNA’s effect on intrinsic fitness is greater on-drug than off-drug (addiction), negative when the effect is greater off-drug than on-drug (expected wt behavior), and zero when the effect is equal on-drug and off-drug (resistance without addiction). We further restricted the inferred set of addicted sgRNAs (*A >* 0) to those that comprised at least 5% of the pool by week 5, indicating robust growth while on-drug (**Figs. 2C-D, S7**). By this metric, the fitness of the CXC-targeting sgRNAs as well as others declined upon drug removal and were identified as addicted sgRNAs (**Supplementary Data S3)**. Altogether, our CRISPR-addiction scan suggests that multiple resistance mutations enriched in the presence of PRC2 inhibitors may in fact act as drug addiction mutations.

Consistent with this idea, we successfully isolated clonal Karpas-422 CXCdel cells by single-cell sorting into media supplemented with 1 μM GSK343. The isolated Karpas-422 clonal cell line contains a heterozygous *EZH2* CXCdel mutation in trans with the Y646N mutation (**Figs. S8, S9A**). To determine if the CXCdel mutation preferentially resides in trans with respect to the activating mutations (Y646N or A682G), we performed amplicon sequencing of the complementary DNA (cDNA) encompassing both mutation positions in the bulk cell populations. CXCdel mutation frequencies were higher in the wt *EZH2* allele in Karpas-422, suggesting some degree of allelic bias against the Y646N mutation (**Fig. S10A-C**). However, little allele preference was observed with respect to the A682G mutation in Pfeiffer, suggesting that genetic differences in DLBCL cell lines may influence allelic preference (**Fig. S10D**).

Karpas-422 CXCdel^+/–^clonal cells exhibited a bell-shaped dose-response curve, indicating a requirement for an optimal concentration of either PRC2 inhibitor for proliferation (**Fig. 2E**). A similar bell-shaped dose-response curve was also observed for resistant Pfeiffer cells transduced with EZH2 sgD597 and maintained with 200 nM GSK343 (**Fig. S9B, S11-S12**,). Cell cycle analysis of Karpas-422 CXCdel^+/–^ revealed an accumulation of sub-G_1_ cells upon withdrawal of GSK343 indicating growth arrest (**Fig. 2F**). These data support the notion that CXCdel mutant lymphoma cells depend on PRC2 inhibitors for survival and are drug addicted.

To investigate how EZH2 CXCdel impacts PRC2 activity, we assessed H3K27me3 levels in wt and CXCdel cells. While wt cells showed reduced H3K27me3 levels under GSK343 treatment, this reduction was attenuated in both Karpas-422 and Pfeiffer CXCdel cells (**Fig. 3A**), suggesting that resistance is mediated by maintaining levels of H3K27me3 despite the presence of PRC2 inhibitors. To assess the impact of the CXCdel mutation independent of the EZH2 hyperactivating mutations found in the lymphoma cell lines, we knocked in the CXCdel mutation into the endogenous wt *EZH2* locus in K562 cells (**Figs. S9C, S12**) and found that H3K27me3 levels were globally increased in homozygous K562 CXCdel^+/+/+^compared to K562 wt cells (**Figs. 3B, S13**). To study EZH2 CXCdel hyperactivity biochemically, we purified both wt and CXCdel recombinant human PRC2 (EZH2 wt or CXCdel, EED, SUZ12, RBAP48, and AEBP2) (**Fig. S14**) (*19*). PRC2 CXCdel exhibited a ten-fold increase in methyltransferase activity, demonstrating that the mutation’s hypermorphic effect is intrinsic to the PRC2 complex itself (**Fig. 3C**). The EZH2 CXC domain mediates the interaction between PRC2 and the substrate nucleosome (*13, 14*), leading us to consider if enhanced PRC2 activity is due to increased substrate affinity. In support, electrophoretic mobility shift assays (EMSA) demonstrated that PRC2 CXCdel has increased affinity towards mononucleosomes (**Figs. 3D, S15**).

**Fig. 3.**
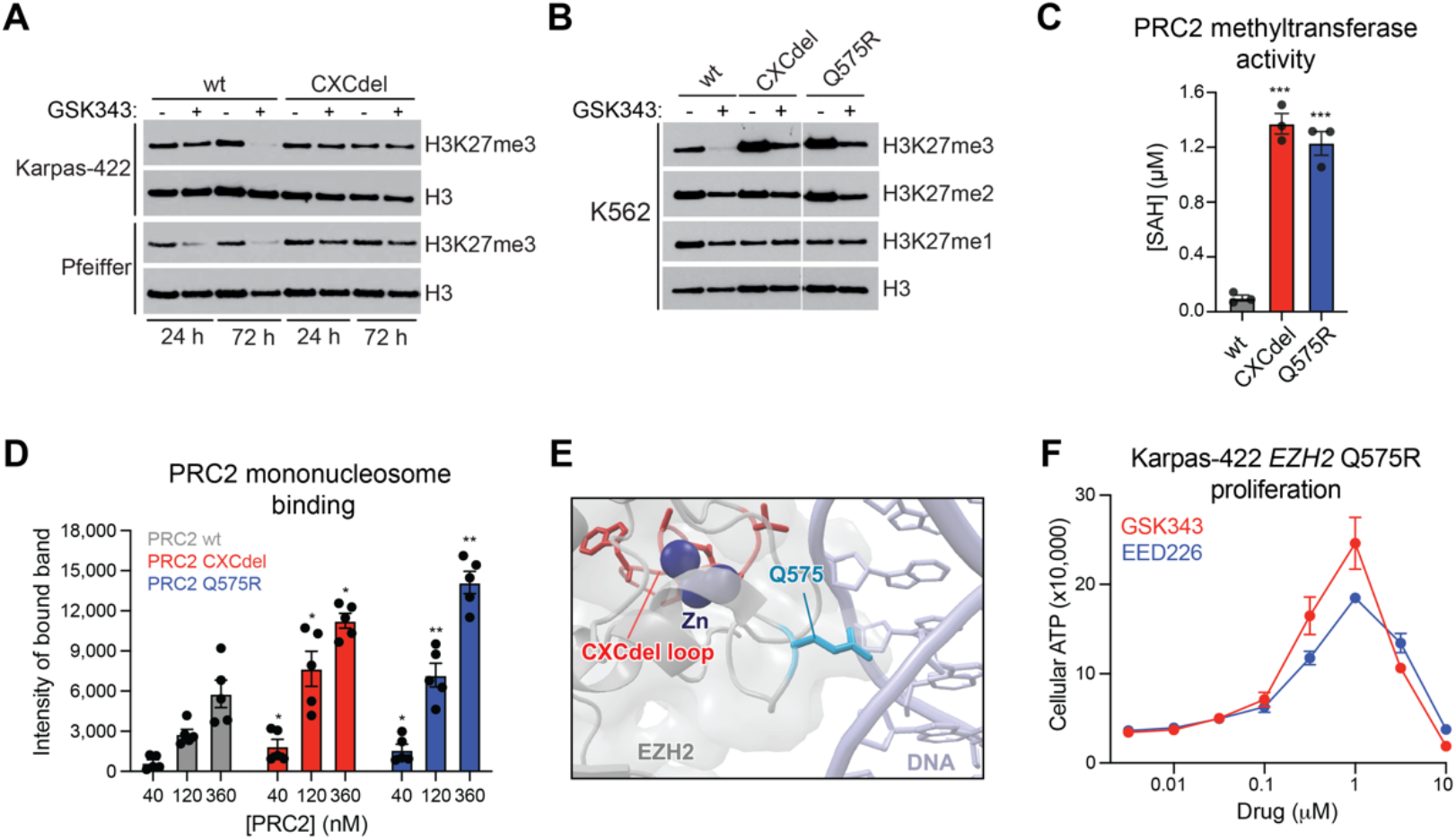
EZH2 CXC-mutants allosterically activate PRC2 methyltransferase activity by increasing substrate affinity. **A**. Immunoblot showing H3K27me3 levels in wt and CXCdel Karpas-422 and Pfeiffer cells after treatment with vehicle, 1 μM GSK343 (Karpas-422) or 200 nM GSK343 (Pfeiffer) for 24 or 72 hours. **B**. Immunoblot showing H3K27me1/2/3 levels in wt, CXCdel, and Q575R K562 cells after treatment with vehicle or 5 μM GSK343 for 72 hours. **C**. Barplot showing enzyme activity of wt, CXCdel, and Q575R PRC2 as measured by production of *S*-adenosyl homocysteine (SAH) by the MTaseGlo Methyltransferase Assay. \*\*\**P* < 0.001 (two-tailed Student’s t-test conducted between wt and mutant PRC2). Bars represent mean ± s.e.m across three replicates. **D**. Quantification of the PRC2-nucleosome bound band across five EMSA replicates to measure binding of wt, CXCdel, and Q575R PRC2 to 185 bp mononucleosomes. \**P* < 0.05 \*\**P* < 0.01 (two-tailed Student’s t-test conducted between wt and mutant PRC2 at each concentration of PRC2). Bars represent mean ± s.e.m across five replicates. **E**. Structural view showing the location of EZH2 Q575 with respect to nucleosomal DNA. The EZH2 CXCdel loop is shown in red, Q575 residue in light blue, and zinc ions are in dark blue. EMDB-7306, PDB: 6WKR **F**. Dose-response proliferation curves of Karpas-422 EZH2 Q575R under GSK343 and EED226 treatment for 10 days. Data represent mean ± s.e.m across three replicates.

While most cancer mutations in the EZH2 CXC domain abrogate PRC2 activity (*20, 21*), the hyperactivity of EZH2 CXCdel suggests that some EZH2 CXC domain mutations might also enhance PRC2 activity by increasing nucleosome binding. Supporting this idea, *EZH1* Q571R, a mutation present in 27% of thyroid adenomas, has been shown to activate PRC2 activity (*22*). The homologous residue in EZH2, Q575, inserts into the major groove of DNA wrapping the substrate nucleosome, suggesting that its mutation to a positively charged residue might increase nucleosome binding (**Fig. 3E**) (*13, 14*). Supporting this notion, purified PRC2 containing EZH2 Q575R exhibited elevated methyltransferase activity and mononucleosome binding relative to wt PRC2 (**Figs. 3C-D, S14, S15**). Moreover, K562 cells containing a knock-in of *EZH2* Q575R also exhibited elevated H3K27 methylation (**Figs. 3B, S9C, S12**). As such, we next considered if EZH2 Q575R confers drug addiction. We knocked in the *EZH2* Q575R mutation into Karpas-422 cells in the presence of 1 μM EED226 (**Figs. S9A, S12**). The *EZH2* Q575R mutant cells displayed biphasic growth responses to increasing concentrations of EED226 and GSK343, demonstrating drug addiction (**Fig. 3F**). Thus, both deletions and point mutations in the EZH2 CXC domain can augment PRC2 nucleosome binding and methyltransferase activity to confer drug addiction in EZH2-mutant lymphoma cells.

To investigate how EZH2 CXCdel hyperactivity impacts the chromatin landscape, we profiled H3K27me3 using quantitative spike-in ChIP-seq (ChIP-Rx) in wt and CXCde^+/–^Karpas-422 cells – continuously grown in GSK343 and after 72 hours of drug withdrawal (*23*). Upon drug withdrawal, H3K27me3 strikingly increased and broadly spread across the genome in CXCdel^+/–^cells (**Figs. 4A-B, S16A-B**), suggesting that H3K27me3 might encroach into neighboring regions typically blocked by active histone modifications (*24*–*26*). To determine which regions are most affected by H3K27me3 spreading upon drug removal, we assessed the correlation between each active histone modification with the change in H3K27me3 signal at the borders of H3K27me3 domains upon drug withdrawal. H3K36me3 was most associated with increases in H3K27me3 (**Figs. 4C, S16C-D**). In agreement, we observed a greater increase in H3K27me3 within the gene bodies compared to promoters of expressed genes for CXCdel^+/–^cells (**Fig. 4D**), consistent with the enrichment of H3K36me3 at gene bodies (**Fig. S16E**) (*27*). H3K27me3 ChIP-Rx also revealed spreading and broad gains in H3K27me3 genome-wide and within gene bodies in CXCdel^+/+/+^versus wt K562 cells (**Fig. S16F-H**). Despite these global changes in H3K27me3 levels induced by EZH2 CXCdel, genome-wide binding of EZH2 was largely unchanged between CXCdel and wt cells for both Karpas-422 and K562 (**Fig. S16I-J**). RNA-seq before and after drug withdrawal in Karpas-422 CXCdel^+/–^cells revealed 338 differentially expressed genes (log_2_(fold-change) > ±0.5, *p*_adj_ < 0.05), of which 218 genes (64.5%) were down-regulated after 72 hours of drug removal (**Fig. 4E, Supplementary Data S6**). For the down-regulated genes, an even greater increase in H3K27me3 within gene bodies was observed upon drug withdrawal (**Fig. S16K**). Of note, several key genes related to B-cell proliferation and survival, including *MYB, BCL9*, and *CDK6*, were down-regulated and gained H3K27me3 within their gene bodies (**Fig. 4A, 4E, Supplementary Data S6**) (*28, 29*). Together, our data suggest that drug withdrawal in CXCdel mutant cells induced pervasive spreading of H3K27me3, which is pronounced at gene bodies marked with H3K36me3, to silence critical genes necessary for lymphoma cell survival.

**Fig. 4.**
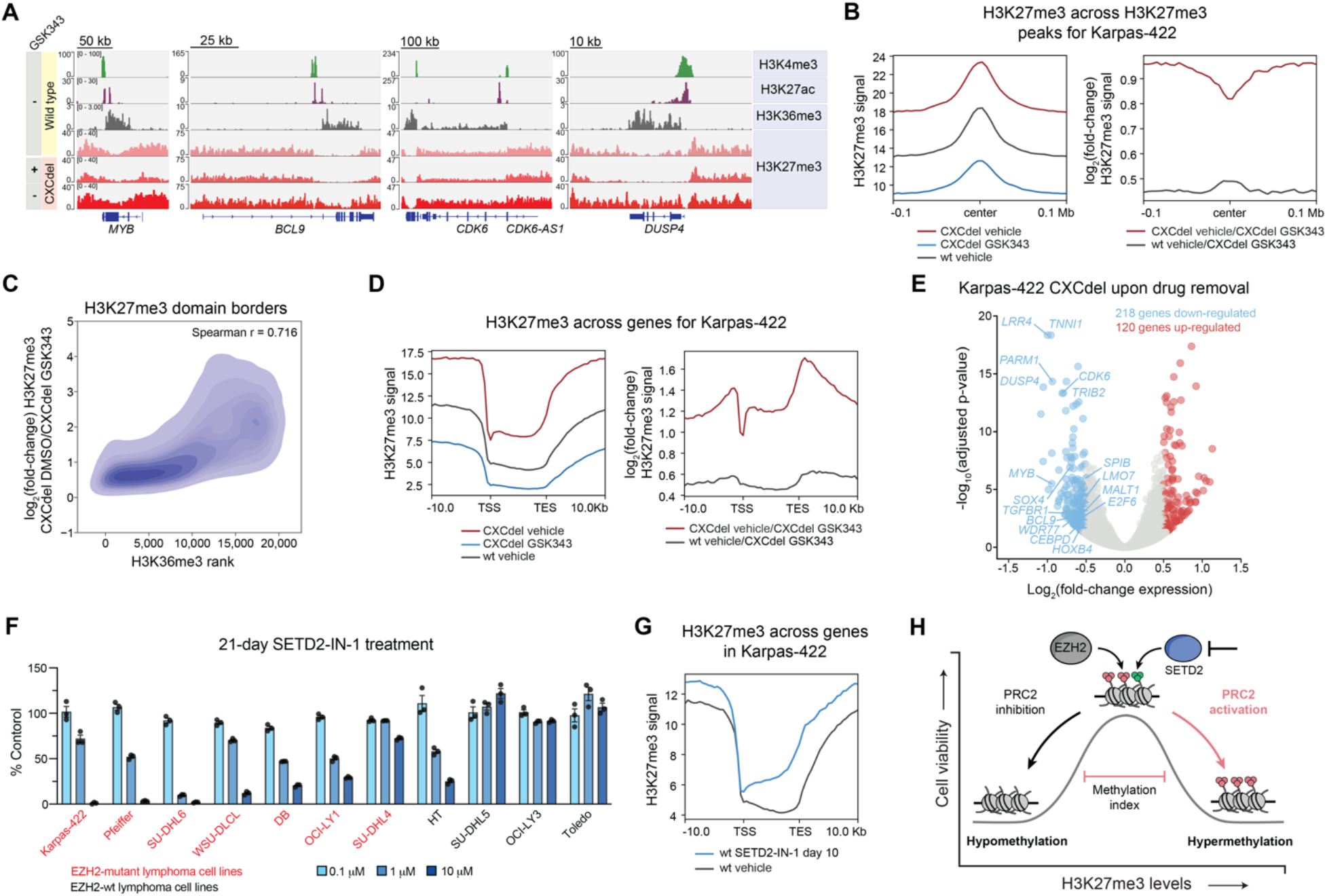
EZH2 CXCdel induces pervasive spreading of H3K27me3 and phenocopies SETD2 inhibition. **A**. Representative genome browser views for histone modification ChIP-seq in wt Karpas-422 and H3K27me3 ChIP-Rx performed in wt and CXCdel^+/–^Karpas-422 cells under continuous GSK343 (1 µM) treatment versus 3 days post drug removal. **B**. Aggregate profile plots of Karpas-422 H3K27me3 ChIP-Rx signal averaged across replicates (*y*-axis, left) and log_2_(fold-change) H3K27me3 ChIP-Rx signal relative to GSK343-treated Karpas-422 CXCdel^+/–^cells averaged across replicates (*y*-axis, right) centered around H3K27me3 domains in wt Karpas-422 cells (*x*-axis). **C**. Density plot depicting log_2_(fold-change) H3K27me3 ChIP-Rx signal between vehicle- and GSK343-treated Karpas-422 CXCdel^+/–^cells (*y*-axis) at wt Karpas-422 H3K27me3 domain borders relative to ChIP-seq signal rank for H3K36me3 (*x*-axis). **D**. Aggregate profile plots of Karpas-422 H3K27me3 ChIP-Rx signal averaged across replicates (*y*-axis, left) and log_2_(fold-change) H3K27me3 ChIP-Rx signal relative to GSK343-treated Karpas-422 CXCdel^+/–^cells averaged across replicates (*y*-axis, right) centered around gene bodies (*x*-axis). (TSS: transcription start site; TES: transcription end site). **E**. Volcano plot showing log_2_(fold-change) in gene expression in Karpas-422 CXCdel^+/–^cells under continuous GSK343 (1 µM) treatment versus 72 hours post drug removal. **F**. Bar plots showing cell proliferation in the presence of SETD2-IN-1 at 0.1, 1, or 10 µM after 21-day treatment. EZH2-mutant lymphoma cell lines are in red and EZH2-wt lymphoma cell lines are in black. Bars represent mean ± s.e.m across three replicates. **G**. Aggregate profile plot of Karpas-422 H3K27me3 ChIP-Rx signal averaged across replicates (*y*-axis) for SETD2 inhibitor conditions centered around gene bodies (*x*-axis). **H**. Model illustrating the repressive methylation index in EZH2-mutant lymphomas.

Our findings demonstrate a biphasic dependency on H3K27me3 levels to maintain EZH2-mutant lymphoma cell proliferation, revealing the potential of activating PRC2 to block lymphoma cell growth. Given the observation that H3K27me3 aberrantly spreads into H3K36 methylated regions, we tested whether inhibition of SETD2, the only known methyltransferase to deposit H3K36me3 and a PRC2 antagonist, could enable H3K27me3 spreading and halt DLBCL proliferation. We treated a panel of EZH2-wt and -mutant lymphoma cell lines with SETD2-IN-1, a recently disclosed, selective SETD2 inhibitor (*30*). Of the seven EZH2-mutant cell lines tested, six were sensitive to SETD2-IN-1 in comparison to only one of the four EZH2-wt cell lines (**Fig. 4F**). Profiling H3K36me3 and H3K27me3 in Karpas-422 cells using ChIP-Rx revealed that SETD2 inhibition substantially decreased levels of H3K36me3 and led to H3K27me3 spreading on chromatin (**Fig. S17A-D, F**). This spreading was pronounced in gene bodies of expressed genes, particularly those down-regulated in Karpas-422 CXCdel^+/–^cells upon GSK343 withdrawal (**Figs. 4G, S17E**). Collectively, these data suggest that SETD2 inhibition selectively blocks the growth of EZH2-mutant DLBCL cell lines, in part by inducing H3K27me3 spreading, nominating the activation of PRC2 as a strategy to block EZH2-mutant lymphoma cell growth.

In summary, through the identification of addiction mutations, we discover that a H3K27me3 ceiling exists in EZH2-mutant lymphoma. This H3K27me3 ceiling forms the upper limit of a “repressive methylation index” – a window of H3K27me3 levels permissible for lymphoma cell proliferation (**Fig. 4H**). Activating EZH2 mutations in lymphoma elevate H3K27me3 levels to block B-cell differentiation but in doing so create a ‘Goldilocks’ epigenetic state near this repressive methylation ceiling. PRC2 activation, which can be achieved by the inhibition of antagonistic factors like SETD2, represents an approach to target EZH2-mutant lymphoma by pushing the cells beyond this ceiling. Our work raises the prospect that identification of drug addiction mechanisms involving mutations in the direct drug target may be broadly exploited for discovery of cancer vulnerabilities. Consequently, explicitly searching for drug addiction mutations using methods like CRISPR-addiction scanning is warranted to interrogate these possibilities across other epigenetic complexes and different drug target classes altogether.

## Acknowledgments

We thank P. Gosavi, members of T. Cech laboratory especially A. Gooding and Y. Long, and members of B. Kingston laboratory especially S. Marr and T. Oei for guidance on protein biochemistry experiments; P. Cole for providing the nucleosomal DNA sequence for binding studies; K. Ngan for guidance on computational analysis of CRISPR-suppressor scans; M. Quezada for aiding in genomics analysis; D. Youmans, D. Narducci, and A. Hansen for guidance on live-cell imaging experiments; R. Ryan for providing cell lines for SETD2 inhibitor studies; the Bauer Core Facility at Harvard University, particularly Zachary Nizioleck and Jeffery Nelson for their assistance with cell sorting. We thank members of the Liau laboratory for helpful discussions and comments on the manuscript.

## Author contributions

H.S.K., A.M.F., and B.B.L. conceived the study and designed experiments; H.S.K. and A.M.F. performed and analyzed cell and molecular biology experiments; A.M.F., A.L.W. and J.W.M performed protein purification and biochemical experiments; H.S.K. performed genomics experiments; H.S.K. and A.P.S. analyzed genomics data; S.M.K. performed growth rate calculations for addicted sgRNAs; H.S.K., A.M.F. and B.B.L. edited and wrote the manuscript, with inputs from all authors; B.B.L. held overall responsibility for the study.

## Data and materials availability

ChIP–seq and RNA-seq data have been deposited to NCBI GEO (GSE199889). Transformed CRISPR-suppressor scanning reads (log_2_ + 1) used for Figs. 1, 2, S1 and S5 are supplied in Supplementary Dataset 2. Addiction scores and growth rates used for Figs. 2 and S7 are supplied in Supplementary Dataset 3. Computer code employed in Fig. 2 for calculation of the addiction score is publicly available at https://github.com/skissler/EZH2 under the GNU General Public License, Version 3.0 (details included in code repository). All other computer code employed in Figs. 1, 2 and 4 are available from the corresponding author upon reasonable request.

## Materials and Methods

### Cell culture and lentivirus production

K562 was obtained from ATCC (CCL-243); 293FT was purchased from Thermo Fisher Scientific (R70007; Karpas-422 and Pfeiffer were gifts from B.E. Bernstein (Dana-Farber Cancer Institute); SU-DHL6, WSU-DLCL, DB, OCI-LY1, SU-DHL4, HT, SU-DHL5, OCI-LY3 and Toledo were gifts from R. Ryan (University of Michigan). SF9 was obtained from Expression Systems (94-001F). All cell lines were authenticated by Short Tandem Repeat profiling (Genetica) and routinely tested for mycoplasma (Sigma-Aldrich). All media for human cell lines were supplemented with 100 U/mL penicillin and 100 µg/mL streptomycin (Gibco) and FBS (Peak Serum). All human cell lines were cultured in a humidified 5% CO2 incubator at 37 °C. K562 and Pfeiffer were cultured in RPMI-1640 (Gibco) supplemented with 10% FBS. Karpas-422 were cultured in RPMI-1640 supplemented with 20% FBS. 293FT cells were cultured in DMEM (Gibco) supplemented with 10% FBS, 0.1 mM MEM Non-Essential Amino Acids (NEAA), 6 mM L-glutamine and 1 mM MEM Sodium Pyruvate. SF9 was cultured in a non-humidified, non-CO2 incubator at 27 °C shaking at 120 rpm in ESF921 media (Expression Systems) supplemented with 50 U/mL penicillin and 50 µg/mL streptomycin. For lentivirus production, plasmids were co-transfected with *GAG/POL* and *VSVG* plasmids into 293FT using FuGENE HD (Promega). Media was exchanged after 7 h and the viral supernatant was collected 48–72 h after transfection and filtered (0.45 µm). Pfeiffer and Karpas-422 were transduced by spinfection at 1,070 x *g* (5 acceleration, 2 brake) for 90 min at 37 °C with 8 µg/ml polybrene (Santa Cruz Biotechnology). After 48 h post-transduction, media was changed and puromycin (Thermo Fisher Scientific) selection was carried out for 7 days at 1 µg/mL for Karpas-422 and Pfeiffer.

### Chemical reagents

Compounds were stored at −80 °C in 100% DMSO. The vehicle condition represents 0.1% DMSO treatment. GSK343 (Cat No. S7164) and EED226 (Cat No. S8496) were purchased from Selleck Chemicals (≥99% purity by HPLC). SETD2-IN-1 (Cat No. HY-136328) was purchased from MedChemExpress (≥99% purity by HPLC).

### sgRNA pooled cloning and CRISPR-suppressor scanning experiments

The *PRC2* tiling library included all sgRNA with an NGG PAM cleavage site and off-target score (MIT Specificity Score) greater than 20 within the coding sequence of the three members of the core PRC2 complex: EZH2 (*NP_004447*.*2*), EED (*NP_001294936*.*1*) and SUZ12 (*NP_056170*.*2*) (*31*). These sgRNA sequences are listed in **Supplementary Data S1**. Oligos containing the sgRNA sequences were cloned into pLentiCRISPR.v2 in a pooled fashion as previously described (*31*). pLentiCRISPR.v2 was a gift from F. Zhang (Addgene plasmid number 52961, Broad Institute). Lentiviral particles carrying the resultant *PRC2* tiling library were generated as described above and titered according to published procedure (*32*). For CRISPR-suppressor scanning, Karpas-422 cells (4.8 × 10^7^) were transduced at a multiplicity of infection < 0.3 and subsequently selected with puromycin. After puromycin selection, cells were split into pools and treated with either 1 µM GSK343, 1 µM EED226 or vehicle. Genomic DNA was isolated at specified time points using the QIAamp® Blood Mini Kit (Qiagen). To measure the composition of the population, sgRNA sequences for all replicates were amplified with PCR primers (**Supplementary Data S4**) and sequenced as previously described (*9*).

### Validation of enriched sgRNAs from CRISPR-suppressor scan

sgRNAs were cloned into pLentiCRISPR.v2 (**Supplementary Data S5**); Karpas-422 and Pfeiffer cells were transduced with the resultant plasmids and selected with puromycin. Transduced Karpas-422 cells were treated with 1 µM of GSK343 or 1 µM of EED226 for 6 weeks. Transduced Pfeiffer cells were treated with incremental concentrations of GSK343 or EED226 (for GSK343 − 50 nM for 2.5 weeks, 100 nM for 1.5 weeks and 200 nM for 2 weeks; for EED226 - 200 Nm EED226 for 2.5 weeks and 500 nM EED226 for 3.5 weeks). Viability was monitored by flow cytometry with Helix NP™ NIR viability dye (BioLegend).

### Generation of clonal drug-resistant EZH2 mutant cell lines and genotype determination

Karpas-422 cells were transduced with pLentiCRISPR.v2 *EZH2* sgD597 and treated with 1 µM GSK343 for 6 weeks to enrich for drug-resistant mutant cells. Surviving cells were single cell-sorted into media containing 1 µM GSK343. Genomic DNA was isolated as mentioned above. For library preparation, genomic PCR primers (**Supplementary Data S4**) with Illumina adapter sequences were used to amplify specified regions of EZH2 as previously described (*9*). Samples were sequenced on a MiSeq genome analyzer (Illumina). The sequencing reads were analyzed using CRISPResso2 (v.2.0.40) (*33*).

### Generation of knock-in cell lines

To generate CXCdel, Q575R K562 and Q575R Karpas-422 knock-in cell lines, electroporation was performed using NEON system with gRNA complex and Alt-R Cas9 enzyme from IDT. gRNA complex was formed by reconstituting gene-specific crRNA (**Supplementary Data S5**) and tracrRNA (1072534, IDT) to 100 µM in IDT duplex buffer, mixed 1:1 and hybridized by incubation at 95°C for 5 min, followed by cooling to room temperature on the bench top. RNP complex was formed by mixing gRNA complex with Alt-R Cas9 enzyme and followed by incubation at room temperature for 15 min. 2 × 10^5^ cells were washed twice with PBS and resuspended in buffer R. RNP complex, gene-specific Ultramer ssODN donor (**Supplementary Data S5**) and Alt-R Cas9 electroporation enhancer were added to the cell suspension and electroporated at 1350V with 10 ms pulse width for 4 pulses. After electroporation, cells were immediately transferred to prewarmed media supplemented with HDR Enhancer (10007910, IDT). To generate single-cell clones, cells were sorted on a MoFlo Astrios EQ cell sorter.

### Cell growth assays

Cell lines were seeded in 96-well plates with 45,000 cells per well in triplicate with drug or vehicle treatments. The number of viable cells was quantified at day 3, day 7, day 10 and day 14 with CellTiter-Glo (Promega) according to manufacturer’s instructions on a SpectraMax i3x plate reader. ATP standard curve was prepared using known concentrations of ATP and used to calculate the ATP content of cells. On the days of measurement, cells were replated with fresh media containing drug or vehicle.

### Cell cycle analysis

Wild-type and CXCdel Karpas-422 and Pfeiffer cells were treated with GSK343 (1 µM for Karpas-422, 200 nM for Pfeiffer) or vehicle for 7 days. Cells were washed with PBS and then fixed with cold 70% ethanol and incubated for a minimum of 2 h at −20°C. After fixing, cells were washed twice with PBS and then resuspended in Propidium Iodide (PI) staining solution (1x PBS supplemented with 50 µg/mL PI, Invitrogen; 100 µg/mL RNase A, Qiagen; and 2 mM MgCl_2_) and incubated for 30 min at room temperature in the dark). Cells were analyzed by flow cytometry (ACEA Novocyte, Agilent). Data was analyzed with FlowJo™ v10 software.

### Immunoblotting

For whole cell extracts, cells were lysed on ice using radioimmunoprecipitation assay (RIPA) buffer (Boston BioProducts) supplemented with fresh HALT™ protease inhibitor cocktail (Thermo Fisher Scientific). The lysate was clarified by centrifugation at 18,000 *x g* for 15 min. For isolation of core histones, lysates were prepared using the Histone Extraction Kit (Active Motif) according to the manufacturer’s protocol. Protein concentration of the lysates was measured using Bradford Assay (Bio-rad). Immunoblotting was performed according to standard procedures. The primary antibodies used are as follows: EZH2 (Cell Signaling Technology, #5246); H3K27me1 (Active Motif, cat no. 61015); H3K27me2 (Cell Signaling Technology); H3K27me3 (Cell Signaling Technology, #9733); H3K36me3 (Cell Signaling Technology, #9763); (Histone 3 (Active Motif, cat no. 39763); GAPDH (Santa Cruz Biotechnology, sc-477724).

### Purification of recombinant PRC2

Human recombinant PRC2 used in this study contained 5 members: EZH2, SUZ12, EED, RBAP48 and AEBP2 (UniProtDB entry isoform sequences Q15910-2, Q921E6-3, O75530-1, Q09028-1, Q6ZN18-1 respectively). Recombinant CXCdel PRC2 contained the same 5 complex members, with a 10 amino acid deletion in EZH2 from T592-S601. Recombinant Q575R PRC2 contained the same 5 complex members. All members were tagged with an N-terminal MBP tag cleavable by PreScission protease. pFastBac plasmids for the five wild-type PRC2 members were gifts from T. Cech. The proteins were co-expressed in insect cells as according to the Bac-to-Bac baculovirus expression system. Detection of gp64 was used to determine baculovirus titer (Expression Systems). For expression, SF9 cells were grown to a density of 2 106 cells/mL and infected with PRC2 baculovirus at a MOI of 1.2 for AEBP2 and 0.7 for all other subunits. The cells were incubated for 72 h (27 °C, 120 rpm), harvested and then frozen with liquid nitrogen for future purification.

PRC2 complexes were purified according to a literature procedure (*19, 34*). All protein purification steps were performed at 4 °C. Insect cells were first lysed by incubating for 1 h with lysis buffer (10 mM Tris-HCl, pH 7.9, 250 mM NaCl, 0.5% Nonidet P-40 Substitute, 1 mM TCEP) at a ratio of 8 mL of buffer per gram of cell pellet. The lysate was clarified by centrifugation at 29,000 *x g* for 40 min. The clarified lysate was added to equilibrated Amylose resin (NEB Cat no. E8021S, 0.3 mL of resin per gram of cell pellet) and bound in batch for 2 h. The resin was then washed with >10 column volumes of lysis buffer, followed by >16 column volumes (CV) of Wash buffer 1 (10 mM Tris-HCl, pH 7.9, 500 mM NaCl, 1 mM TCEP) and lastly >16 CV Wash buffer 2 (10 mM Tris-HCl, pH 7.9, 150 mM NaCl, 1 mM TCEP). PRC2 was eluted by addition of 3 CV of Wash buffer 2 + 10 mM Maltose. Protein was concentrated to 15 mg/mL using a Amicon Ultra-15 Centrifugal Filter Unit, 30 kDa molecular weight cut off (MWCO). In-house made PreScission protease was added at a ratio of 1:100 (protease:PRC2) and the NaCl concentration was adjust to 250 mM by addition of the appropriate amount of 5 M NaCl. After overnight incubation, cleavage of the MBP tags was confirmed by SDS-PAGE. Next, the cleaved protein was subjected to a 5mL HiTrap Heparin column at a flow rate of 1.5 mL/min with a gradient elution from Buffer A (10 mM Tris-HCl, pH 7.9, 150 mM NaCl, 1 mM TCEP) to Buffer B (10 mM Tris-HCl, pH 7.9, 2 M NaCl, 1 mM TCEP) over 35 CV. The salt concentration was diluted to 350 mM and the protein was concentrated and subjected to size-exclusion over a Superose 6 Increase 10/300 GL column with running buffer (20 mM HEPES, pH 7.9, 150 mM NaCl, 1 mM TCEP) at a flow rate of 0.5 mL/min. PRC2-peak fractions were verified with SDS-PAGE. The correct fractions were pooled and concentrated as indicated above. The final concentration was determined by UV absorbance at 280 nm measured by Nanodrop. The ratio of absorbance at 260 nm/280 nm ratio was less than 0.7 indicating minimal nucleic acid contamination.

### Purification of mononucleosomes

*Xenopus* histones were expressed in *E. coli* and purified as previously described with minor modifications as noted here (*35, 36*). Briefly for the histone purification, inclusion bodies were prepared and extracted as described previously (*35, 36*). These pellets were subsequently dissolved into unfolding buffer (7 M guanidinium HCl, 20 mM Tris-HCl, pH 7.5, 10 mM DTT), dialyzed into SAU-200 buffer (7 M Urea, 20 mM sodium acetate pH 5.2, 200 mM NaCl, 5 mM 2-mercaptoethanol, 1 mM EDTA) and loaded onto a 5mL HiTrap SP HP column, eluted by gradient elution over 30 CV from SAU-200 buffer to SAU-600 buffer (7 M Urea, 20 mM sodium acetate pH 5.2, 600 mM NaCl, 5 mM 2-mercaptoethanol, 1 mM EDTA). Histone-containing fractions were pooled, dialyzed into distilled water containing 2 mM 2-mercaptoethanol (at least 3 changes) and lyophilized for storage at −20 °C (*37*). For histone octamer assembly, the histones were first dissolved in in unfolding buffer (20 mM Tris-HCl pH 7.5, 6 M guanidinium HCl, 5 mM DTT), combined in equimolar amounts and dialyzed into refolding buffer (10 mM Tris-HCl pH 7.5, 2 M NaCl, 1 mM EDTA, 5 mM b-ME) and purified over a Superdex 200 column (GE Healthcare). Octamer-containing fractions were pooled according to purity and histone stoichiometry, and then flash-frozen for storage at −80 ºC after addition of glycerol to a final concentration of 10% (v/v).

The 185-base pair (bp) nucleosome DNA used for all studies with *Xenopus* nucleosomes was atcgctgttcaatacatgc<u>acaggatgtatatatctgacacgtgcctggagactagggagtaatccccttggcggttaaaacgcgggg gacagcgcgtacgtgcgtttaagcggtgctagagctgtctacgaccaattgagcggcctcggcaccgggattctccag</u>ggcggccgc gtatagggat. The 601-nucleosome positioning sequence is underlined. The plasmid containing the 185-bp nucleosomal DNA sequence was a gift from Philip Cole. The 185-bp nucleosomal DNA sequence was prepared via large scale plasmid preparation followed by cleavage with the EcoRV HF restriction enzyme. The cleavage product was purified with Q Sepharose HP resin with a stepwise elution from Buffer A (20 mM Tris pH 8.0, 1 mM EDTA) to 40% Buffer B (20 mM Tris pH 8.0, 1 mM EDTA, and 1 M NaCl) and elution with 70% Buffer B. Eluted DNA was subsequently purified by size-exclusion on a Superdex 200 column in Buffer B. The resulting product was buffer-exchanged into Buffer A and stored at −20 °C. Nucleosomes were reconstituted by mixing the DNA and the octamer in a 1:1.1 ratio and performing gradient dialysis as described previously (*36*). Reconstituted nucleosomes were purified by anion exchange with a TSKgel DEAE-5PW (7.5 mm x 7.5 cm) column; their composition was confirmed by native gel electrophoresis and were subsequently concentrated and stored as described previously (*36*). All dialysis steps employed 6-8 kDa MWCO dialysis tubing (Spectra-Por) and 10 kDa MWCO Amicon centrifugal filter units were used for concentration and desalting.

### Electrophoretic mobility shift assay with mononucleosomes

45 nM of mononucleosomes purified as above were incubated with increasing amounts of protein in 20 μL of assay buffer (50 mM Tris-HCl, pH 7.5, at 25 °C, 25 mM KCl, 2.5 mM MgCl_2_, 0.1 mM ZnCl2, 2 mM 2-mercaptoethanol, 0.1mg/mL BSA, 10% v/v glycerol). After incubation at 30 °C for 30 min, 15 μL of the reaction was loaded onto a 3.5% TBE polyacrylamide gel. Gel electrophoresis was performed at 140 V for 1 h in cold 1 × TBE (4 °C). The gel was stained with SYBR Green I (Thermo Fisher Scientific) and fluorescence images were acquired by an Azure c200 imager. Densitometry of the bound PRC2-mononucleosome band was performed by ImageJ. The experiment was repeated five times.

### MTase-Glo Enzyme assay

Endpoint methyltransferase measurements were performed in a 20 μL reaction volume utilizing 80 nM of PRC2 enzyme with 1 μM of mononucleosomes and 40 μM SAM in reaction buffer (50 mM Tris-HCl, pH 8 at 30 °C, 10 mM KCl, 2.5 mM MgCl_2_, 0.1 mM ZnCl_2,_ 2 mM 2-mercaptoethanol, 0.1 mg/mL BSA, 5% v/v glycerol). Reactions were incubated at 30 °C for 2 h and quenched with 0.1 % TFA. Enzyme activity was assessed by SAH production as determined by luminescence through the MTase-Glo assay (Promega). Briefly, SAH production for each reaction was calculated by correlating the raw luminescence to a SAH standard curve correcting for basal luminescence of mononucleosome and buffer to determine SAH production. Each reaction was done in triplicate and the entire experiment was repeated twice.

### ChIP-seq

ChIP was performed as previously described in duplicate on Karpas-422 and K562 cells (*9*). The antibodies used are as follows: EZH2 (Cell signaling Technology, #5246); H3K27me3 (Cell signaling Technology, #9733S); H3K36me3 (Cell signaling Technology, #9763). For quantitative ChIP, drosophila chromatin (Active Motif cat no. 08221011) and spike-in antibody (Active Motif cat no. AB27377370) were added to sonicated chromatin (*23*). Libraries were sequenced on Novaseq sequencer for 100 cycles in paired-end mode.

### RNA-seq library preparation

Total RNA was isolated from cells in triplicate separate cultures using RNeasy Mini Kit (Qiagen). Library preparation was performed using the QuantSeq 3’ mRNA-Seq Library Prep Kit FWD (Lucigen) according to the manufacturer’s protocol. Libraries were sequenced on Novaseq sequencer for 100 cycles in paired-end mode.

### ChIP-seq data analysis

For quantitative ChIP-seq analysis, reads were aligned to a composite reference genome for dm6 and hg38 utilizing bowtie2 (version 2.3.2) according to the Spiker analysis framework (version 1.03). Aligned reads were sorted using samtools (version 0.1.19) and the Spiker split_bam.py script was run to calculate scaling factors and to split reads into an alignment file containing reads specifically aligning to the hg38 genome (Spiker, version 1.0.3) (*38*). This alignment file was then converted into spike-in normalized bigwig file for further analysis and visualization utilizing the deepTools bamCoverage command with the following parameters: -binSize 50 –ignoreDuplicates--normalizeUsing RPKM --ignoreForNormalization chrM --extendReads –scaleFactor. Peaks for H3K27me3 and H3K36me3 were called using Sicer with the parameters: -e 8000 -s hg38 -g 12000 -w 4000 (version 2.0). Individual replicate peaks were subsequently combined utilizing Bedtools2 merge -d 12000 (version 2.26.0). Border regions of peaks were called by using the Bedtools2 “flank” command with the parameter -b 50000 (version 2.26.0). For bin level analysis, the H3K27me3 signal value was averaged across 50 kb bins utilizing the multiBigwigSummary function (deepTools, version 3.4.3). To generate metaprofile plots comparing between treatment conditions the log2 option in the bigwigCompare function was used prior to plotting with the plotHeatmap function (deepTools, version 3.4.3). To generate fingerprint plots investigating H3K27me3 spreading, the plotFingerprint function was used on the hg38 split alignment files for individual replicates with the parameters: --ignoreDuplicates --extendReads --skipZeros (deepTools, version 3.4.3). Subsequent data processing was performed in Python (version 3.9.6) using standard plotting functions.

For EZH2 ChIP-seq analysis, reads were aligned to the hg38 genome utilizing bowtie2 (version 2.3.2). Aligned reads were sorted using samtools (version 0.1.19). This alignment file was then converted into bigwig format for further analysis and visualization utilizing the deepTools bamCoverage command with the following parameters: -binSize 50 --ignoreDuplicates --normalizeUsing RPKM --ignoreForNormalization chrM –extendReads --blackListFileName. Specifically, the file “ENCFF356LFX.bed” downloaded from Encode was used to define blacklisted regions for hg38. For bin level analysis, the EZH2 signal value was averaged across 50 kb bins utilizing the multiBigwigSummary function (deepTools, version 3.4.3). Subsequent data processing was performed in Python (version 3.9.6) using standard plotting functions.

### RNA-seq data processing and differential gene expression analysis

RNA-seq data was processed according to the QuantSeq 3′ mRNA-Seq Library Prep recommended analysis pipeline through alignment to the Ensembl transcriptome (GRCh38). DESeq2 with R (v.3.5.1) was employed for the differential expression analysis. All genes with both an adjusted *P* < 0.05, calculated using the Benjamini–Hochberg correction, and |log_2_(fold-change)| > 0.5 were considered to be differentially expressed. The location of the transcription start site (TSS) and transcription end sites (TES) of genes was determined using the biomaRt package (version 2.41.1) and only genes with a defined hgnc symbol name were included in the annotation list. When multiple isoforms were present, the TSS and TES were defined to be at the minimum and maximum positions, respectively. A table of significantly differentially expressed genes is provided in **Supplementary Data S6**.

### Statistical methods

Statistical parameters including the exact value and definition of *n*, the definition of center, dispersion, precision measures (mean ± S.D. or S.E.M) and statistical significance are reported in figures and figure legends.

## Supplementary Text

### Motivation for the addiction score

In pooled ecological competition experiments, the hallmark of drug addiction is when a lineage’s intrinsic fitness is greater when on drug than when off drug (*i*.*e*., the intrinsic fitness declines when drug is removed). Thus, addiction may manifest for a given lineage as a faster growth rate when on drug than when off drug. However, other processes can also cause a lineage to grow faster on drug than off drug, such as out-competition by more fit strains when drug is removed (*i*.*e*., the relative fitness declines when drug is removed, despite no change in intrinsic fitness). To identify lineages that may be addicted to drug, we therefore require a way to distinguish changes in intrinsic fitness from changes in relative fitness. We can accomplish this by modeling each lineage’s proliferation over time using the laws of competitive logistic growth and comparing each lineage’s growth against a reference lineage that is unaffected by the presence of drug (a “purely resistant” lineage). This approach yields an “addiction score”, defined as the difference between a given lineage’s intrinsic fitness when on drug *vs*. off drug, so that positive values indicate lineages that may be addicted to drug. The following sections outline the calculation of the addiction score.

### Derivation of the addiction score

#### Dynamics of the overall population

Consider a population of cells which consists of many sub-populations (sgRNAs), each of which may have a different intrinsic fitness. We can begin by describing the size *N* of the total population.

Let us assume that the population’s growth is governed by logistic dynamics: when the population size is small, the population exhibits exponential growth, but when the population approaches the carrying capacity *K*, the growth plateaus. These dynamics are described by the standard logistic equation (*18*):

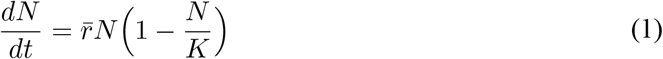

Here, 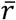 is the overall fitness of the population, *i*.*e*., the exponential growth rate (offspring per unit time) of the population when growth is uninhibited.

To simplify, we can consider the dynamics of the population density, *ρ* = *N/K*, which can take values between 0 and 1. Substituting this expression for *ρ* into Equation 1 gives

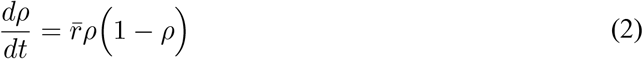

#### Dynamics of individual sgRNAs

Now, let us consider the individual sgRNAs. Let *y*_*i*_ represent the density of sgRNA *i*, so that

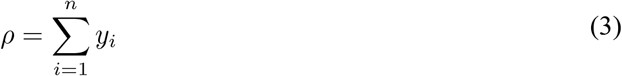

Substituting this into Equation 2 gives

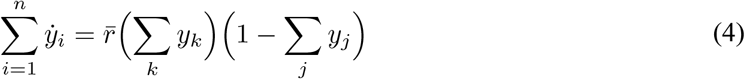

where 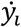 is the derivative of *y*_*i*_ with respect to time. Next, we assume that 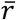 is equal to the mean fitness across all sgRNAs:

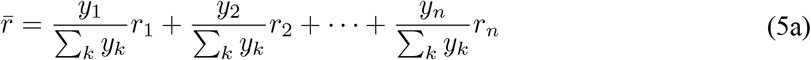

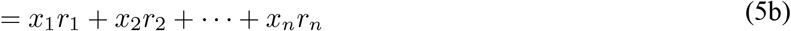

Here, *r*_*i*_ is the intrinsic fitness (uninhibited exponential growth rate) of sgRNA *i* and *x*_*i*_ is the fraction of the population made up by sgRNA *i*.

Next, we seek an equation to describe the growth of an individual sgRNA *i*. We substitute Equation 5a into Equation 4 and express the right-hand side as a sum over *i*:

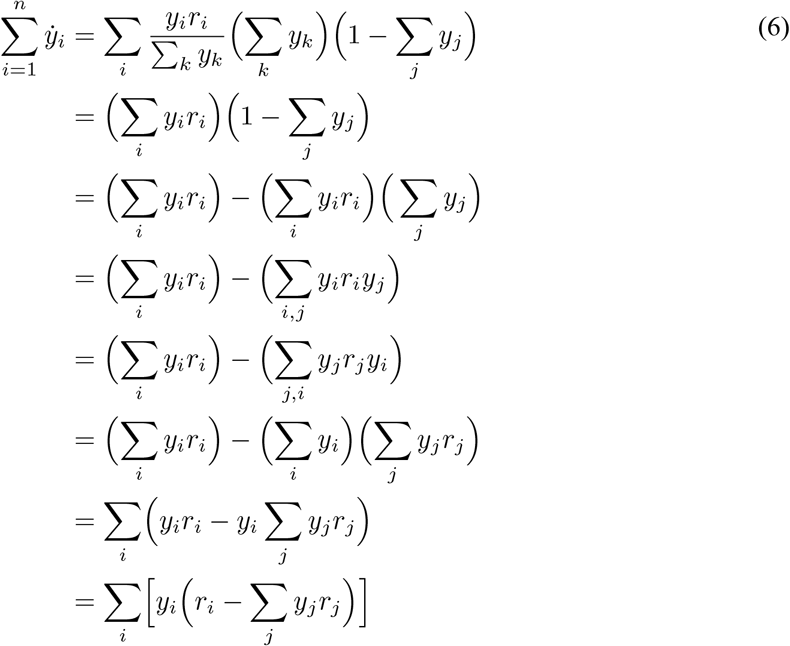

This relation shows that we obtain the correct overall population dynamics if we assume that each sgRNA’s growth is governed by another logistic equation:

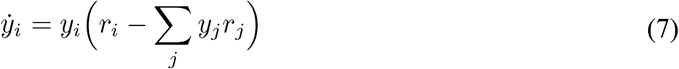

Equation 7 defines a system of equations that is equivalent to the competitive Lotka-Volterra equations (*18*). An interpretation of this equation is as follows: when the overall population size (across all sgRNAs) is small, the growth rate of sgRNA *i* is equal to *r*_*i*_. When the overall population size grows, competition is governed by the term *y*_*i*_ *y*_*j*_ *r*_j_; that is, interactions between sgRNAs *i* and *j* are deleterious to sgRNA *i* by a factor that’s proportional to the fitness of sgRNA *j*.

To illustrate the dynamics resulting from Equation 7, we can numerically simulate the densities of three hypothetical sgRNAs, each starting at a density of 1/500, where drug is applied on days 0 through 8 and removed for days 8 through 16 (**Fig. S18A**).

The intrinsic fitnesses of the sgRNAs while on drug are *r*_*1*_ = 1.51, *r*_*2*_ = 1.50, and *r*_*3*_ = 1.49. When drug is removed, the intrinsic fitnesses of sgRNAs 1 and 2 remain unchanged, but the intrinsic fitness of sgRNA 3 decreases to 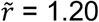 (a change of −0.29).

#### Dynamics of sgRNA proportions

Now, we derive a system of equations for the relative sgRNA proportions. Recall that the relative proportion of sgRNA *i, x*_*i*_, is given by

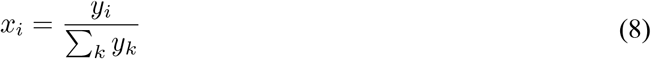

Bearing in mind the relations for *ρ* and 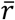 in Equations 3 and 5, the relative proportion of the pool made up by sgRNA *i* over time is described by:

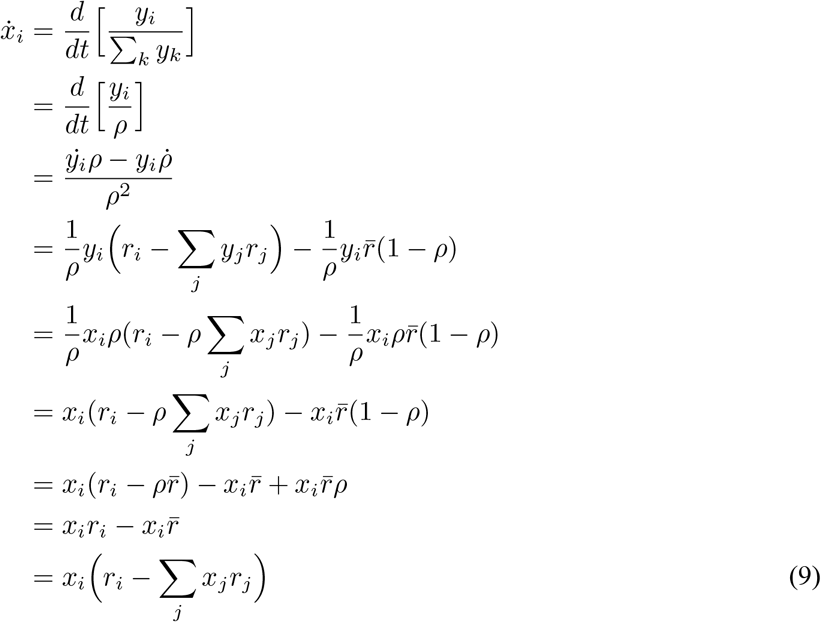

Note that this equation has the same form as Equation 7, which describes the relative densities of the sgRNAs. The only difference is in the initial conditions: in the previous example, we started with *y*_*1*_(0) = *y*_*2*_(0) = *y*_*3*_(0) = 1/500, implying that *x*_*1*_(0) = *x*_*2*_(0) = *x*_*3*_(0) = 1/3. To verify that this is true, we can plot each sgRNA’s density *y*_*i*_ divided by the total population density using Equation 7 (**Fig. S18B**) and compare with the relative frequencies *x*_*i*_ given by Equation 9 (**Fig. S18C**), noting their equality.

#### Calculating change in intrinsic fitness

The next task is to determine the change in a sgRNA’s intrinsic fitness when off drug *vs*. on drug given the relative frequencies of the sgRNAs at (a) time 0 (“start”), (b) when drug is removed (“switch”), and (c) at the end of the experiment (“end”). Let us consider the logarithm of the relative proportions of sgRNAs *i* and *j*:

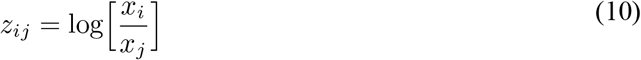

The change in *z*_*ij*_ over time takes a simple mathematical form:

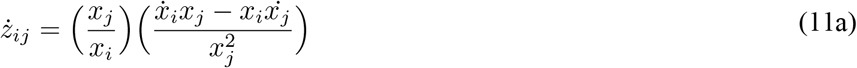

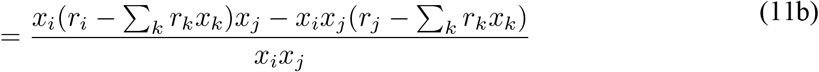

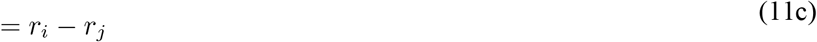

This means that the solution for *z*_*ij*_ is a straight line with slope *r*_*i*_ – *r*_*j*_. For the sgRNAs in our previous example, the dynamics of *z*_*21*_ and *z*_*31*_ are shown in **Fig. S18D**.

Given the sgRNA proportions at the start, switch, and end timepoints, it is straightforward to calculate the slopes of these lines. **Table S1** gives the proportions, *z-*values, and slopes (*σ*_*it*_) for the simulated sgRNAs relative to sgRNA 1.

Next, we can derive an expression for the change in fitness off-drug *vs*. on-drug for each sgRNA. Let us denote the on-drug fitness of sgRNA *i* by *r*_*i*_ and the off-drug fitness of sgRNA *i* by 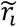. The difference in on-drug *vs*. off-drug fitness is thus 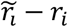. Let us furthermore assume that the fitness of some sgRNA *ξ* (the reference sgRNA) does not depend on drug, *i*.*e*., 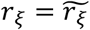. Then, we seek

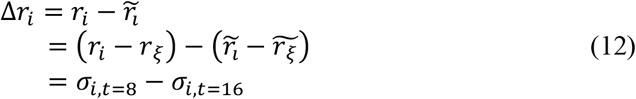

For simulated sgRNA 2 relative to sgRNA 1, the slope at time 8 (*σ*_*2,t=8*_) is equal to the slope at time 16 (*σ*_*2,t=16*_), so 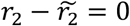; that is, sgRNA 2 is equally fit off-drug as on-drug, which agrees with the construction of the simulation.

For simulated sgRNA 3 relative to sgRNA 1, the slope at time 8 (*σ*_*3,t=8*_) differs from the slope at time 16 (*σ*_*3,t=16*_) by 0.29 units, *i*.*e*., 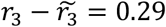. Recall that the on-drug fitness of sgRNA 3 was 1.49 and the off-drug fitness was 1.20, so this also agrees with the construction of the simulation and indicates that simulated sgRNA 3 is addicted to drug.

In general, when Δ*r*_*i*_ < 0, the sgRNA is more fit off-drug than on-drug (wild-type behavior); when Δ*r*_*i*_ = 0, the sgRNA is equally fit off-drug and on-drug (pure resistance); and when Δ*r*_*i*_ > 0, the sgRNA is more fit on-drug than off-drug (addiction). This satisfies the characteristics of an addiction score, so we define the addiction score *A* as

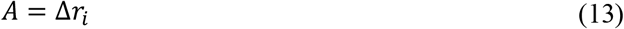

#### Recipe for calculating the addiction score

Taking all of this together, we can calculate change in on-drug *vs*. off-drug fitness if:

- We know the relative frequencies *x* of the sgRNAs at time 0 (*t*_*0*_), when drug is removed (*t*_*rem*_), and when the experiment ends (*t*_*end*_).
- We assume that the overall population is undergoing logistic growth and standard ecological competition.
- We assume that one sgRNA (sgRNA *ξ*) is agnostic to drug (*i*.*e*., has a “pure” resistance mutation).

With these conditions in place, the intrinsic fitness of sgRNA *i* on-drug *vs*. off-drug (the addiction score) is

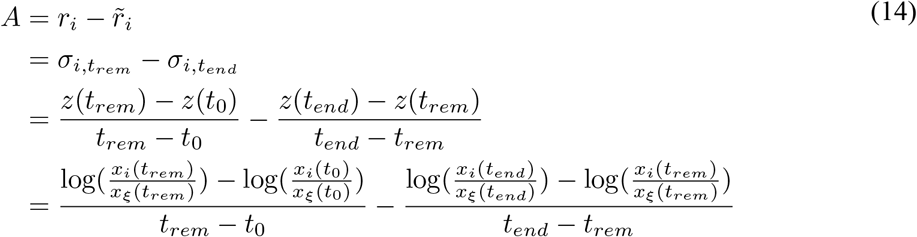

This recipe for calculating the addiction score is implemented in the **Supplementary Code**[https://github.com/skissler/EZH2].

### Simulating the logistic dynamics of the real sgRNAs

To interpolate the relative sgRNA proportions over time given the measured proportions at the start, switch, and end times, we first calculated *r*_*i*_ and 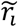 for each sgRNA *i*. We assigned the intrinsic fitness of the reference sgRNA to an arbitrary value (0 for the figures in the main plot, with sensitivity analysis between −10 and 10). In general, the choice of intrinsic fitness for the reference sgRNA may change how quickly the sgRNAs reach a quasi-steady state but does not affect the sgRNA proportions at the start, switch, and end times. In practice, we found very little difference in the dynamics for different assumptions of the intrinsic fitness of the reference sgRNA. We then used the lsoda function from the deSolve package in R version 4.1.3 to numerically solve the system of differential equations given by Equation 9 (see **Supplementary Code** [https://github.com/skissler/EZH2]).

---

**Fig. S1.**
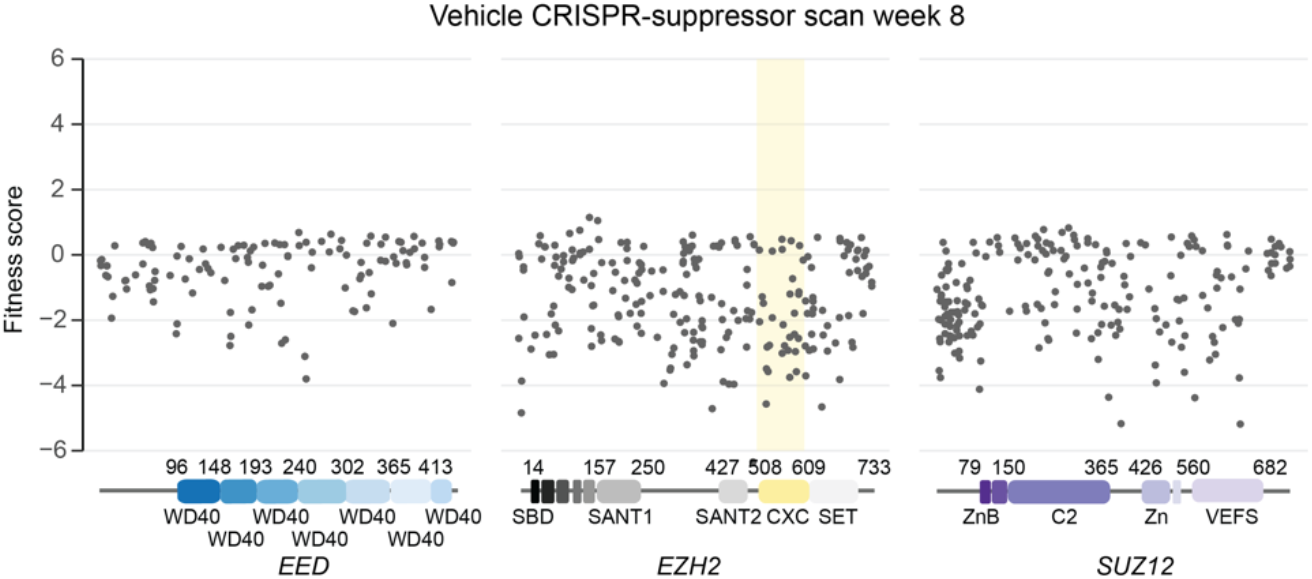
sgRNAs targeting CXC domain or drug binding sites are not enriched in the absence of PRC2 inhibitors. Scatter plots showing fitness scores (*y-*axis) in Karpas-422 under vehicle treatment at week 8. Fitness scores were calculated as the log_2_(fold-change) sgRNA enrichment under vehicle normalized to the mean of the negative control sgRNAs (*n* = 58). The PRC2-targeting guides (*n* = 650) are arrayed by amino acid position in the EED, EZH2 or SUZ12 coding sequence on the *x*-axis corresponding to the position of the predicted cut site. When the sgRNA cut site falls between two amino acids, both amino acids are denoted. Data points represent mean value across three replicates. Protein domains are demarcated by colored panels along the *x*-axis. The location of the EZH2 CXC domain is shown by a yellow background.

**Fig. S2.**
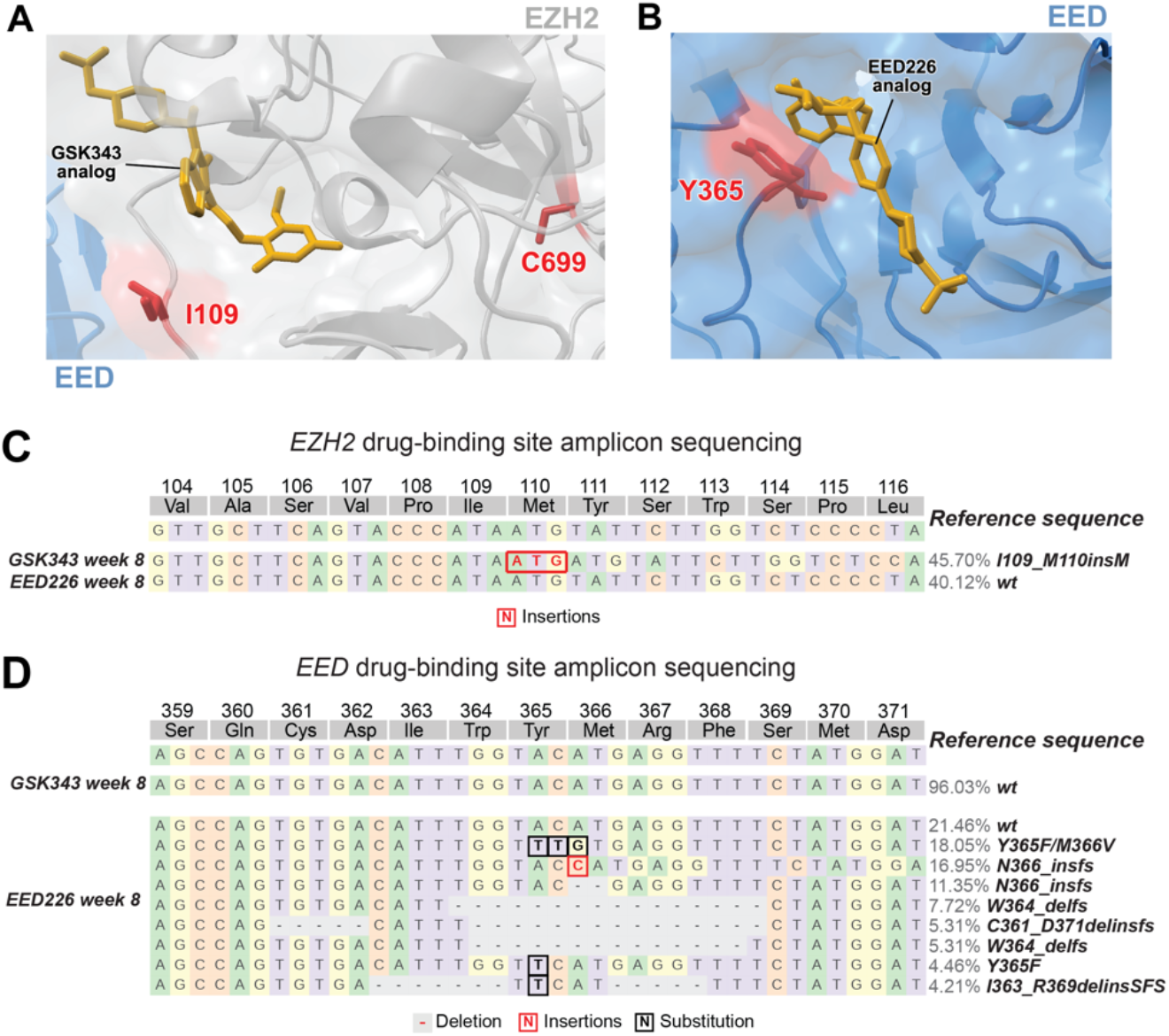
Genotypes enriched in the drug binding pockets of EZH2 and EED observed in the CRISPR-suppressor scan. **A**. Structural view of the GSK343 binding site with EZH2 residues I109 and C699 highlighted in red. GSK343 analog is shown in gold. PDB: 5LS6 **B**. Structural view of the EED226 binding site with EED residue Y365 highlighted in red. EED226 analog is shown in gold. PDB: 5K0M **C**. Schematic showing genotypes and allele frequencies for mutations that are observed at frequencies of > 2% at week 8 in the gDNA encoding *EZH2* SAL domain in Karpas-422 cells in the GSK343 or EED226 CRISPR-suppressor scanning experiments. Shown is a representative out of three replicates. **D**. Same as in **C** but showing the gDNA encoding *EED* WD40 domain.

**Fig. S3.**
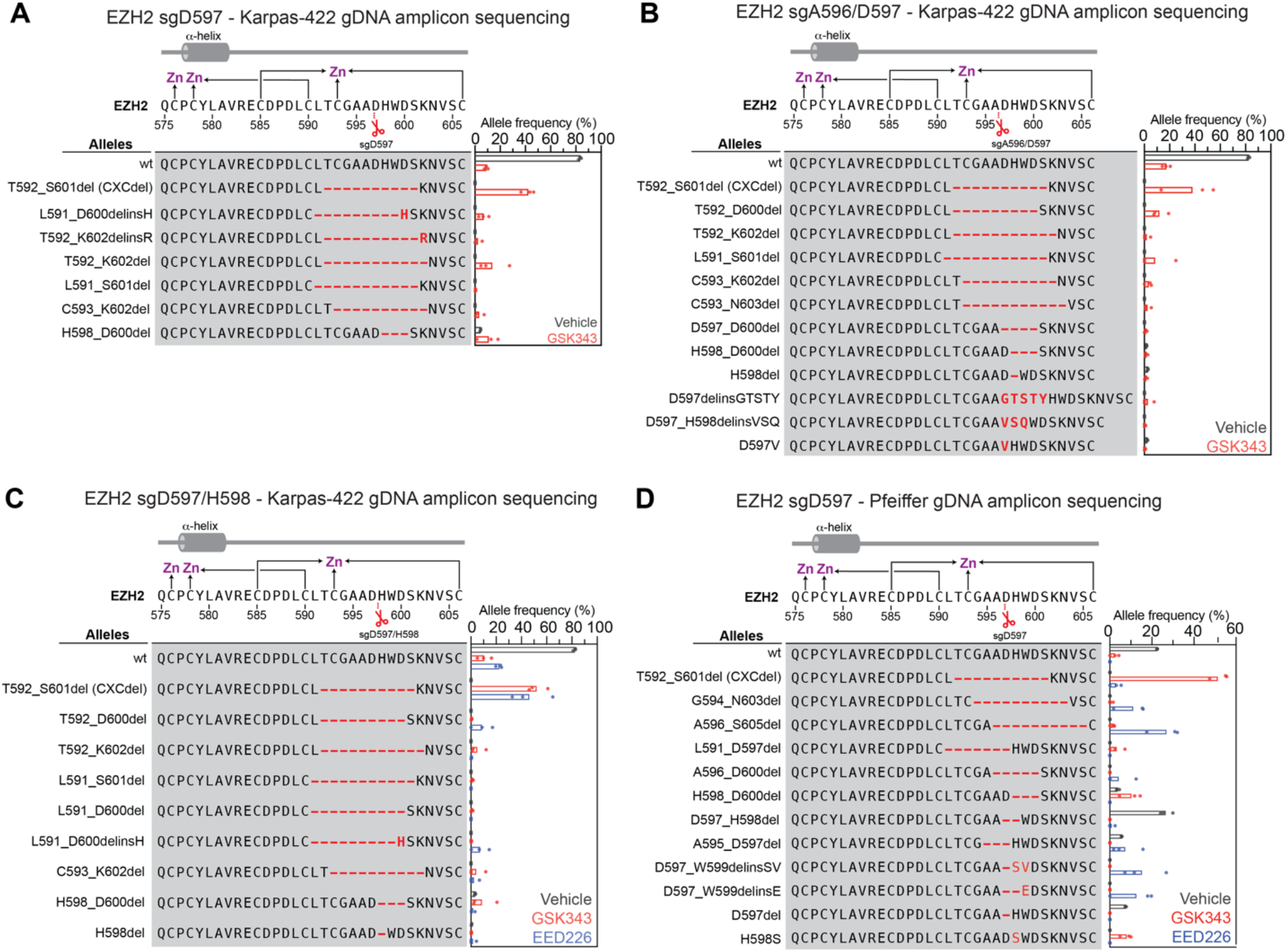
Genotypes enriched upon mutagenesis by CXC-targeting sgRNAs and PRC2 inhibition. **A**. Schematic showing genotypes and bar plots of allele frequencies for mutations that are observed at frequencies of >2% in the gDNA encoding *EZH2* surrounding the CXCdel loop for 6-week GSK343 treatment after transduction with sgD597 in Karpas-422. Allele frequencies under vehicle (grey), GSK343 (red) or EED226 treatment (blue). (top) Schematic depicts the secondary structure of C-terminal CXC domain and cysteine residues that coordinate the zinc ions. **B**. Same as in **A** but for sgA596/D597 transduction in Karpas-422. **C**. Same as in **A** but for sgD597/H598 transduction in Karpas-422. **D**. Same as in **A** but for sgD597 transduction in Pfeiffer cells.

**Fig. S4.**
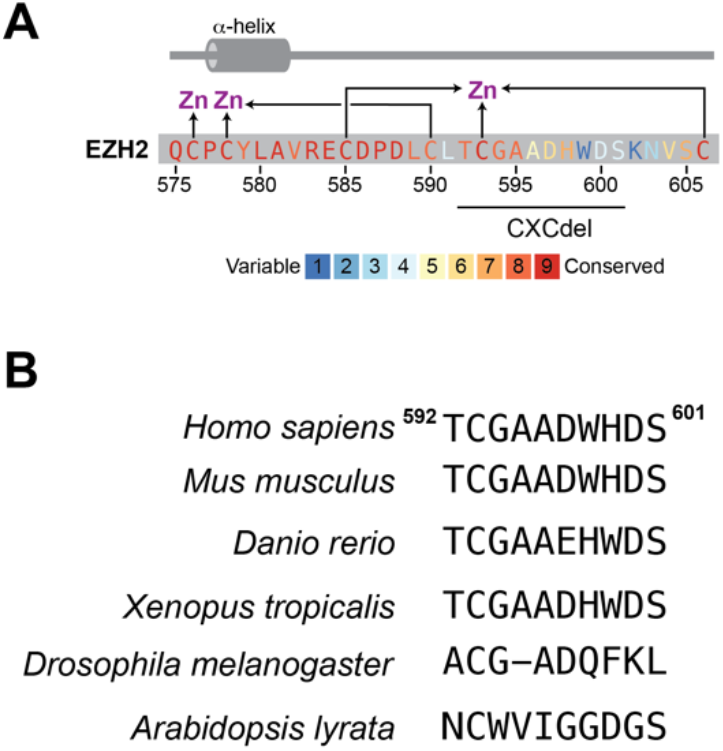
The EZH2 CXCdel loop is poorly conserved. **A**. The conservation scores of each amino acid within and surrounding the EZH2 CXCdel loop are shown in colors ranging from variable (blue) to conserved (red) as calculated by ConSurf-DB (*15*). (top) Schematic depicts the secondary structure of C-terminal CXC domain and cysteine residues that coordinate the zinc ions. **B**. A comparison of amino acids homologous to the EZH2 CXCdel loop in various model organisms.

**Fig. S5.**
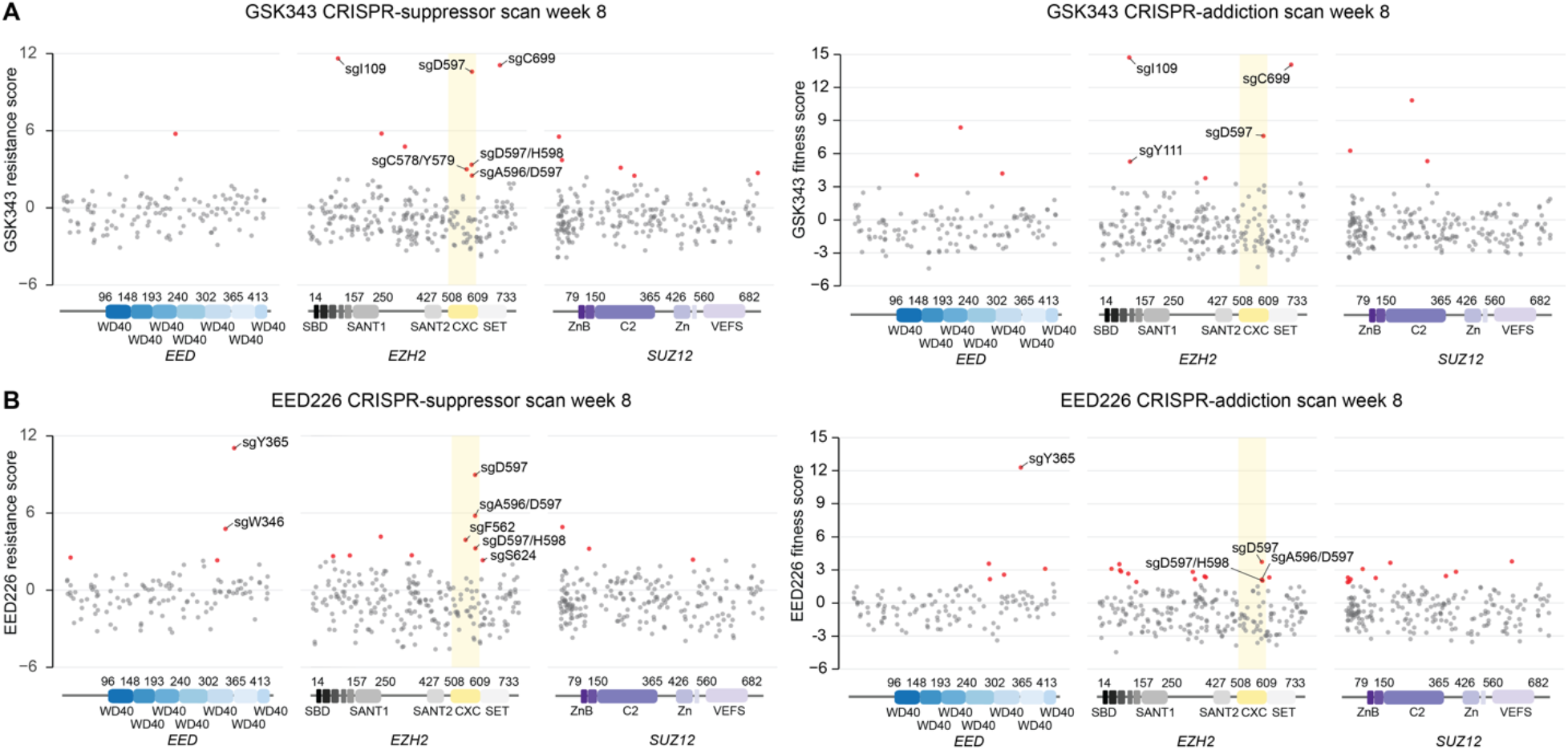
sgRNAs targeting the EZH2 CXC domain are depleted upon drug removal. **A**. Scatter plots showing resistance scores (*y-*axis) in Karpas-422 following 8 weeks of continuous GSK343 treatment (left) and 5 weeks of GSK343 treatment followed by 3 weeks of drug removal (right). Resistance or fitness scores were calculated as the log_2_(fold-change) sgRNA enrichment under drug normalized to the mean of the negative control sgRNAs (*n* = 58). The PRC2-targeting sgRNAs (*n* = 650) are arrayed by amino acid position in the EED, EZH2 or SUZ12 coding sequence on the *x*-axis corresponding to the position of the predicted cut site. When the sgRNA cut site falls between two amino acids, both amino acids are denoted. Data points represent mean value across three replicates. Protein domains are demarcated by colored panels along the *x*-axis. The location of the EZH2 CXC domain is shown by a yellow background. Points colored in red had resistance scores greater than 2 standard deviations (s.d.) above the mean of the negative control sgRNAs. **B**. Same as in **A** but for Karpas-422 following 8 weeks of continuous EED226 treatment (left) and 5 weeks of EED226 treatment followed by 3 weeks of drug removal (right).

**Fig. S6.**
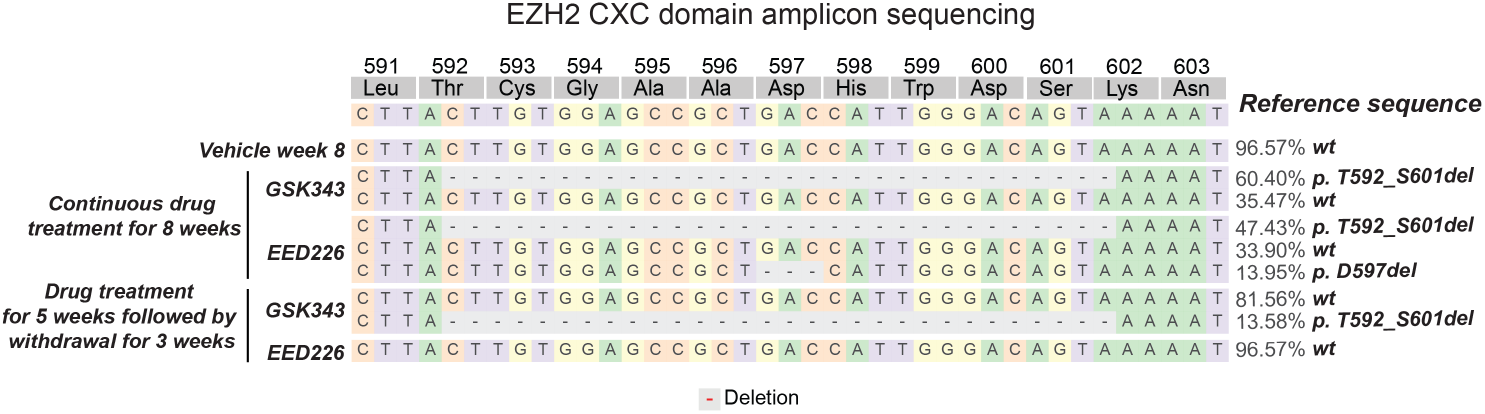
*EZH2* CXCdel is depleted upon drug withdrawal in the CRISPR-addiction scan. Schematic showing genotypes and allele frequencies for mutations that are observed at frequencies of >2% in the gDNA encoding *EZH2* surrounding the CXCdel loop at week 8 of the GSK343 or EED226 CRISPR-addiction scanning experiment. Conditions of the experiment are continuous drug treatment for 8 weeks, continuous vehicle treatment for 8 weeks or 5 weeks of drug treatment followed by 3 weeks of drug withdrawal. All samples were sequenced at the week 8 timepoint. Shown here is a representative out of three replicates.

**Fig. S7.**
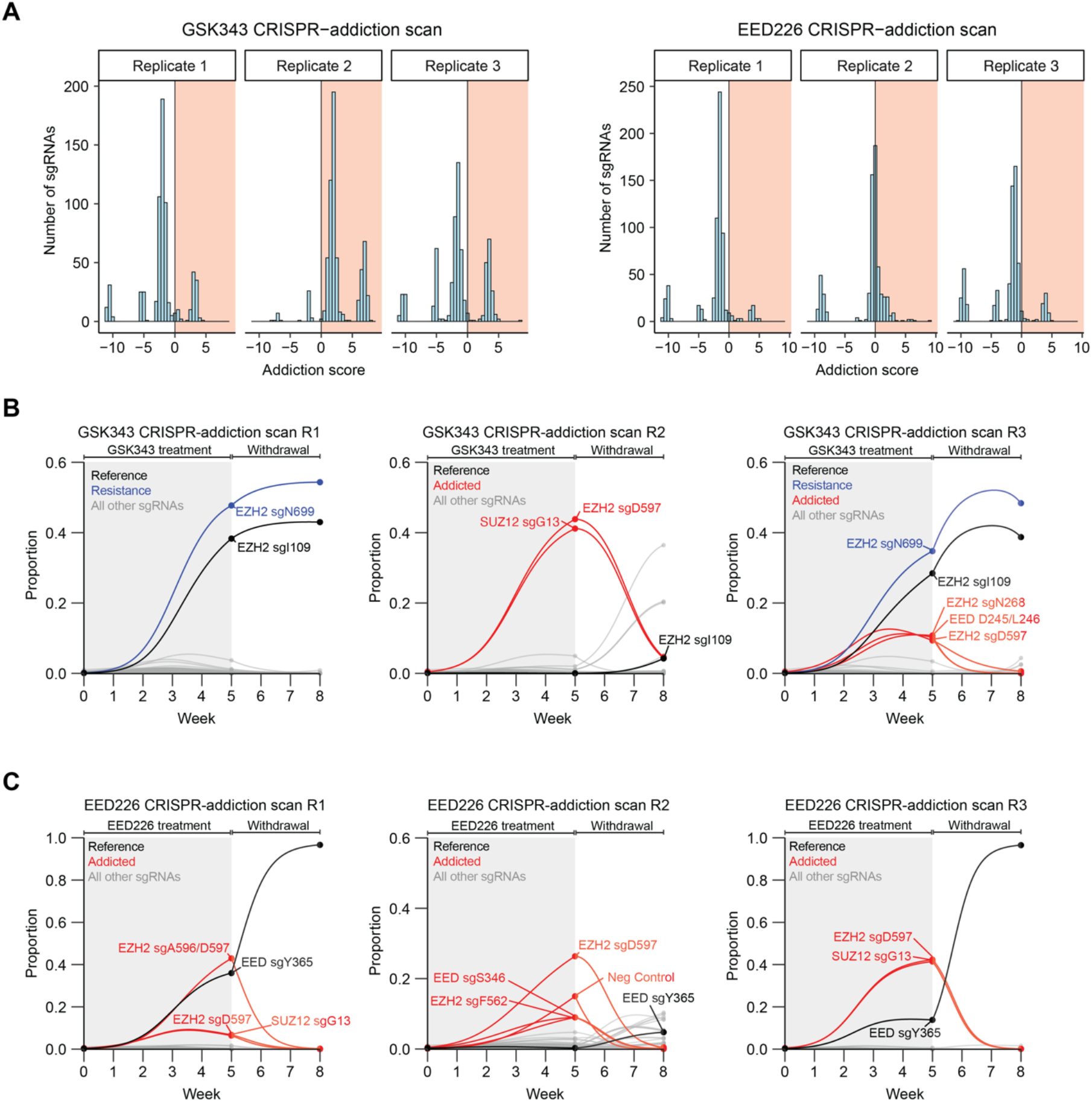
Identification of sgRNAs that induce drug addiction from CRISPR-addiction scanning. **A**. Histogram showing distribution of addiction scores among the three replicates of the GSK343 and EED226 CRISPR-addiction scanning experiments. **B**. Graph showing proportions of each sgRNA (*y*-axis) in each replicate of the GSK343 CRISPR-addiction scan over time (*x*-axis). The grey background indicates the presence of GSK343 and white background indicates the period of drug removal. Curves were calculated using the estimated intrinsic growth rate *r* for each sgRNA-containing cell population under the assumption of competitive logistic growth. The reference sgRNA is shown in black, sgRNAs demonstrating pure resistance are in blue and addicted sgRNAs are in red. **C**. Same as in **B** but for each replicate of the EED226 CRISPR-addiction scan.

**Fig. S8.**
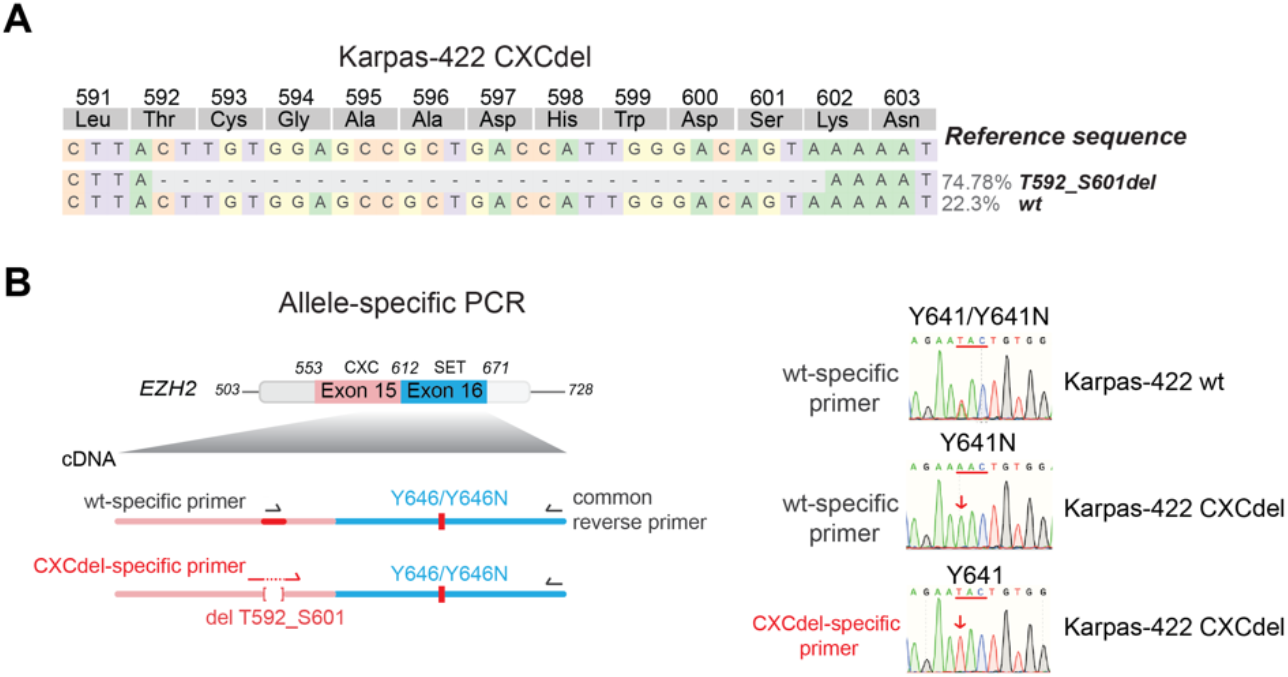
Karpas-422 CXCdel clonal line is heterozygous and resides in trans with Y646N mutation. **A**. Schematic showing genotypes and allele frequencies of the Karpas-422 CXCdel clonal line. We believe that the over-representation of the CXCdel allele in the clonal cell line can be attributed to PCR bias. The heterozygous nature of Karpas-422 CXCdel is confirmed in **B**. **B**. Allele-specific PCR followed by Sanger sequencing to determine if the CXCdel mutation resides in cis or in trans with respect to the Y646N mutation in the Karpas-422 CXCdel clonal cell line.

**Fig. S9.**
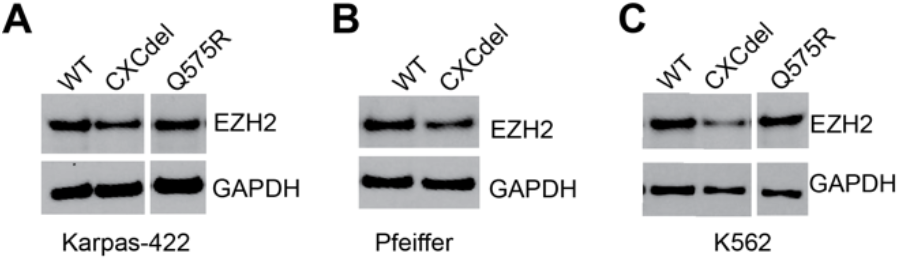
EZH2 expression levels in CXC-mutant cell lines. **A**. Immunoblot showing EZH2 levels in wt, CXCdel, and Q575R Karpas-422. **B**. Same as **A** but for wt and CXCdel mutant Pfeiffer. **C**. Same as **A** but for wt, CXCdel and Q575R K562.

**Fig. S10.**
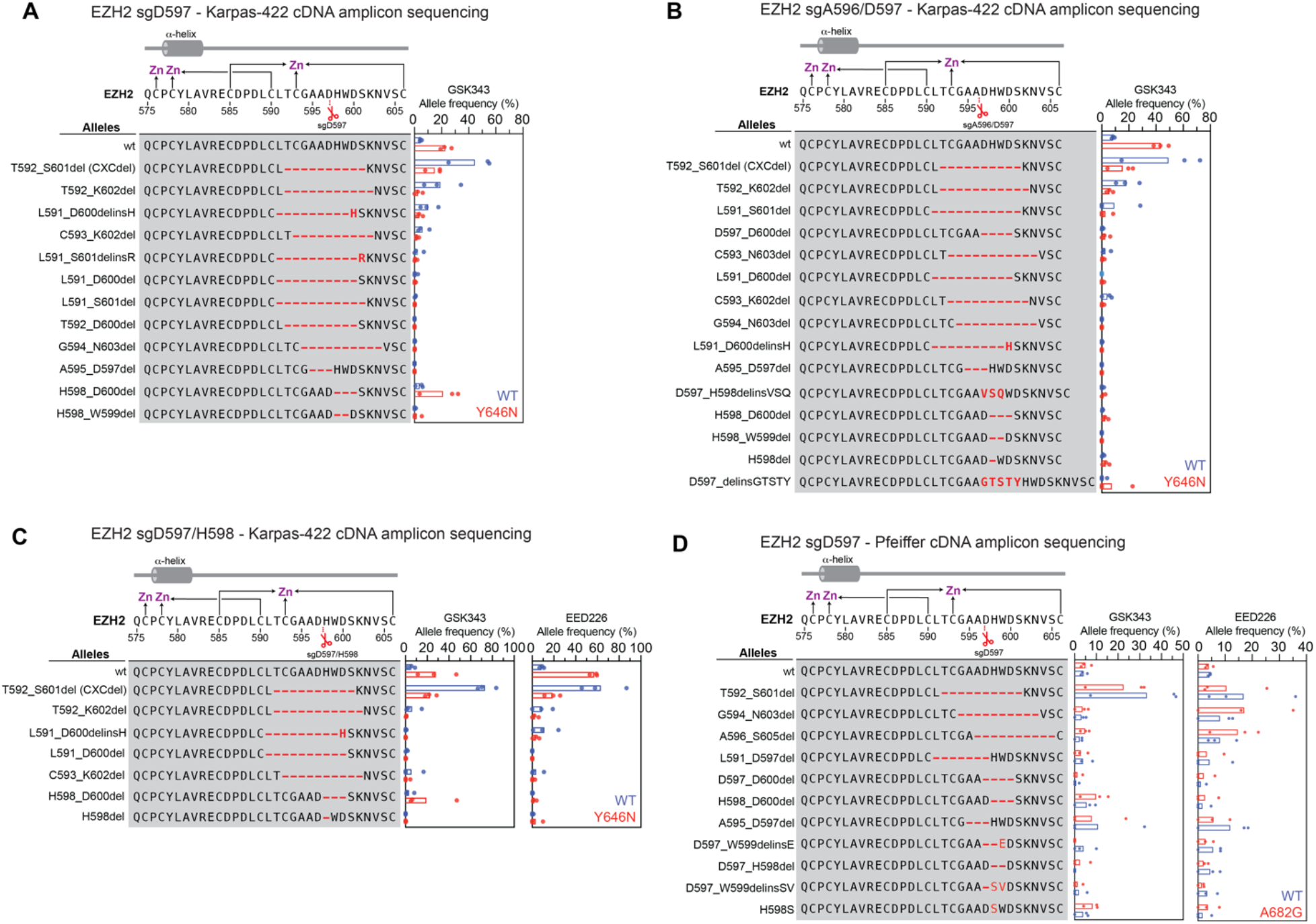
Analysis of allele preference with amplicon sequencing of cDNA in bulk cell population mutagenized by CXC-targeting sgRNAs under PRC2 inhibition. **A**. Schematic showing genotypes and bar plots of allele frequencies for mutations that are observed at frequencies of >2% in the cDNA encoding *EZH2* encompassing the CXC and SET domains for GSK343 treatment at week 6 for sgD597 transduction in Karpas-422. Allele frequencies in the (blue) wt allele and (red) Y646N allele. (top) Schematic depicts the secondary structure of C-terminal CXC domain and cysteine residues that coordinate the zinc ions. **B**. Same as in **A** but for sgA596/D597 transduction in Karpas-422. **C**. Same as in **A** but for sgD597/H598 transduction in Karpas-422. **D**. Same as in **A** but for sgD597 transduction in Pfeiffer. The wt allele is shown in blue and the A682G allele is shown in red.

**Fig. S11.**
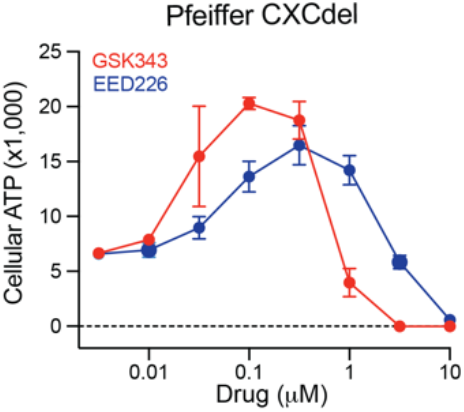
CXCdel Pfeiffer is addicted to PRC2 inhibitors. Dose-response proliferation curves of CXCdel Pfeiffer after treatment with GSK343 and EED226 for 10 days.

**Fig. S12.**
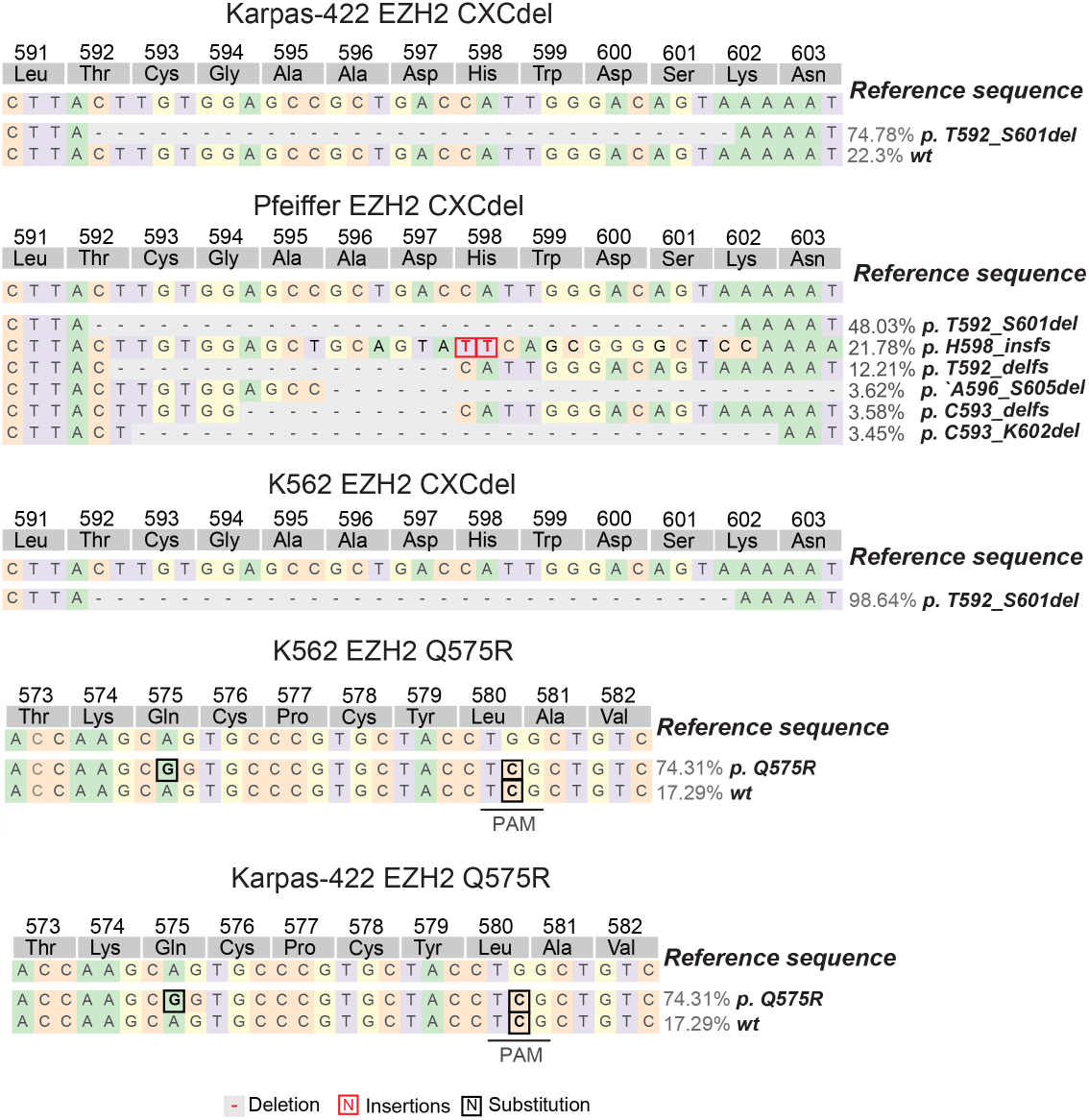
Genotypes of Karpas-422, Pfeiffer and K562 CXC mutants used in this study. Schematic showing genotypes and allele frequencies for mutations that are observed at frequencies of > 2% in the gDNA encoding the EZH2 CXC domain.

**Fig. S13.**
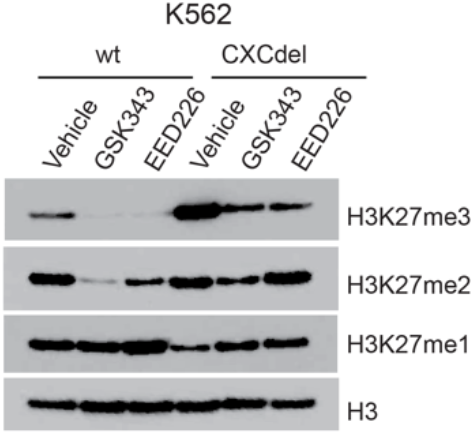
CXCdel mutation leads to global increase in H3K27me3 levels. Immunoblot showing H3K27me1/2/3 levels in wt and CXCdel K562 treated with 5 μM GSK343 or EED226 for 72 h.

**Fig. S14.**
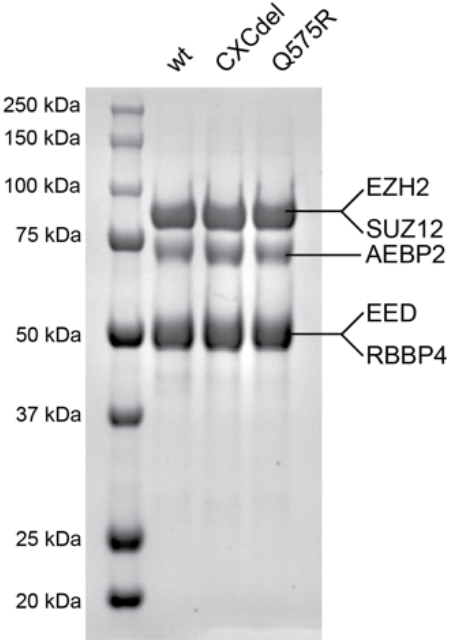
Purification of recombinant 5-member PRC2 complex. SDS-PAGE stained with Coomassie blue showing purified wt, CXCdel, and Q575R PRC2 complexes.

**Fig. S15.**
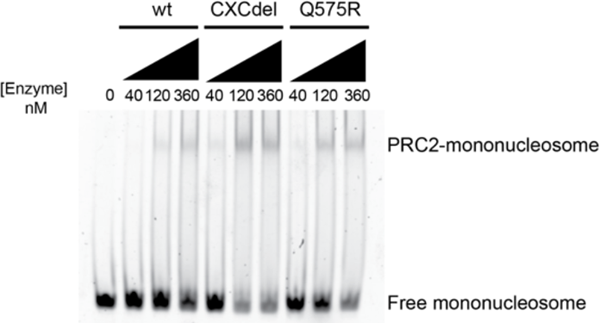
CXCdel and Q575R PRC2 have increased mononucleosome binding relative to wt PRC2. Representative EMSA showing binding of wt, CXCdel and Q575R PRC2 to 185 bp mononucleosomes. The upper band corresponds to the PRC2-mononucleosome complex and bottom band corresponds to the free, unbound mononucleosome.

**Fig. S16.**
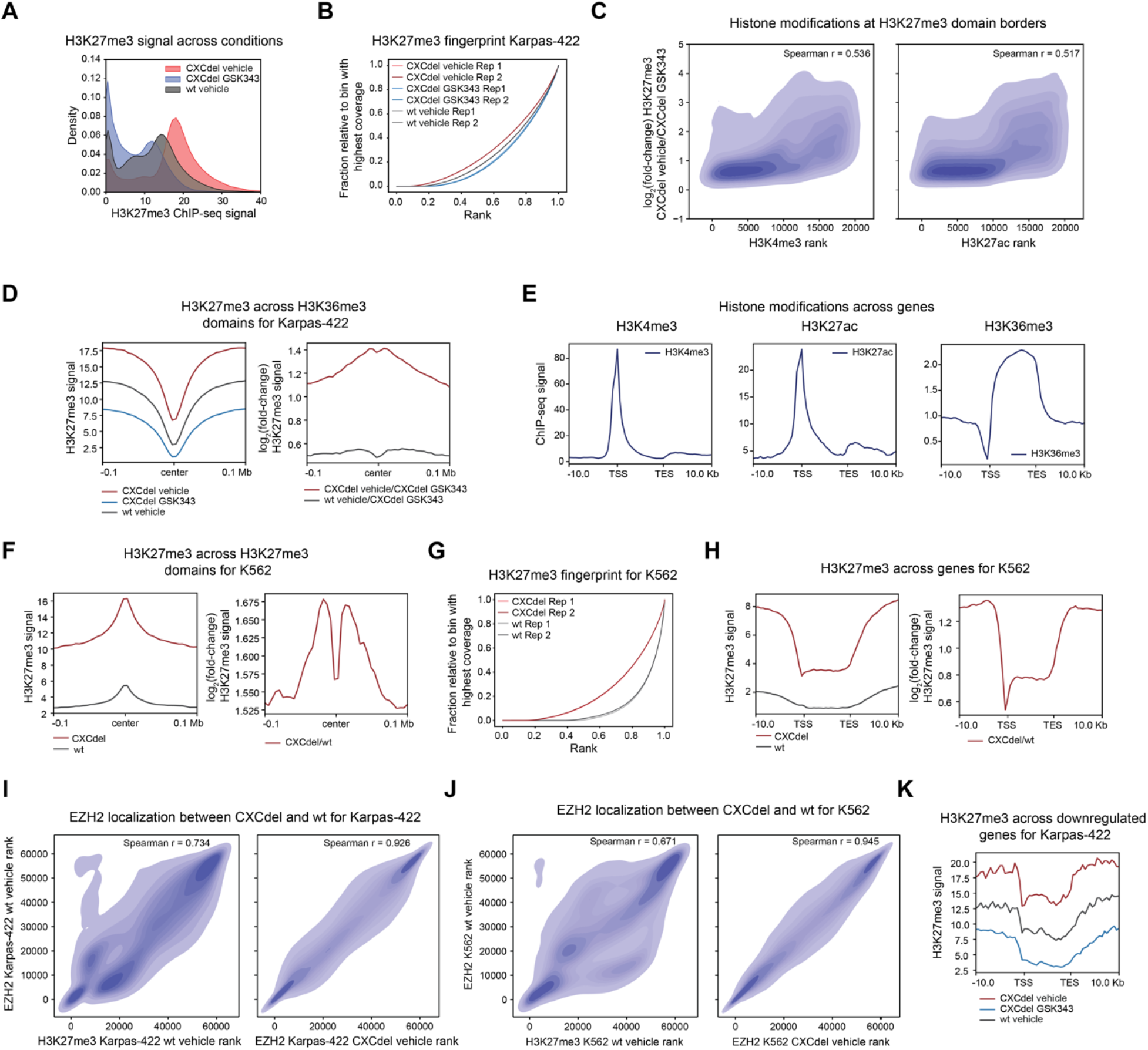
CXC mutation induces pervasive spreading of H3K27me3 in Karpas-422 and K562 cells. **A**. Density plot depicting the distribution of H3K27me3 ChIP-Rx signal averaged across replicates (*x*-axis) of vehicle-treated Karpas-422 CXCdel cells, GSK343 (1 μM) -treated Karpas-422 CXCdel cells, and vehicle-treated wt Karpas-422 cells for 50 kb genomic bins. **B**. Line plot depicting fraction H3K27me3 ChIP-seq coverage relative to that of the bin with the highest coverage (*y*-axis) as a function of the region’s relative genomic rank (*x*-axis) for wt and CXCdel mutant Karpas-422 cells. **C**. Density plot depicting log_2_(fold-change) H3K27me3 ChIP-Rx signal between vehicle-treated CXCdel Karpas-422 cells and GSK343-treated CXCdel cells (*y*-axis) at wt Karpas-422 H3K27me3 domain borders relative to ChIP-seq signal rank for H3K4me3 (left) and H3K27ac (right) (*x*-axis). H3K4me3 rank was calculated from ENCODE data ENCFF932BQZ; H3K27ac rank was calculate from ENCODE data ENCFF435PXJ. **D**. Aggregate profile plots of Karpas-422 H3K27me3 ChIP-Rx signal averaged across replicates (*y*-axis, left) and log_2_(fold-change) H3K27me3 ChIP-Rx signal relative to GSK343-treated Karpas-422 CXCdel cells averaged across replicates (*y*-axis, right) centered around H3K36me3 domains in wt Karpas-422 cells (*x*-axis). **E**. Aggregate profile plots of Karpas-422 ENCODE ChIP-seq signal (*y*-axis) for the specified histone modification centered around gene bodies (*x*-axis). (TSS: transcription start site; TES: transcription end site). H3K4me3 signal was determined from ENCODE data ENCFF932BQZ; H3K27ac signal was determined from ENCODE data ENCFF435PXJ; H3K36me3 signal was determined from ENCODE data ENCFF603BHK. **F**. Aggregate profile plots of K562 H3K27me3 ChIP-Rx signal averaged across replicates (*y*-axis, left) and log_2_(fold-change) H3K27me3 ChIP-Rx signal relative to wt K562 cells averaged across replicates (*y*-axis, right) centered around H3K27me3 domains in wt K562 cells (*x*-axis). **G**. Line plot depicting fraction H3K27me3 ChIP-seq coverage relative to that of the bin with the highest coverage (*y*-axis) as a function of the region’s relative genomic rank (*x*-axis) for wt and CXCdel mutant K562 cells. **H**. Aggregate profile plots of K562 H3K27me3 ChIP-Rx signal averaged across replicates (*y*-axis, left) and log_2_(fold-change) H3K27me3 ChIP-Rx signal relative to wt and CXCdel mutant K562 cells averaged across replicates (*y*-axis, right) centered around gene bodies (*x*-axis). **I**. Density plot depicting EZH2 ChIP-seq signal rank for vehicle-treated wt Karpas-422 cells (*y*-axis) relative to H3K27me3 ChIP-Rx signal rank for vehicle-treated wt Karpas-422 cells (*x*-axis, left) and EZH2 ChIP-seq signal rank for vehicle-treated Karpas-422 CXCdel cells (*x*-axis, right). **J**. Density plot depicting EZH2 ChIP-seq signal rank for wt K562 cells (*y*-axis) relative to H3K27me3 ChIP-Rx signal rank for wt K562 cells (*x*-axis, left) and EZH2 ChIP-seq signal rank for K562 cells CXCdel (*x*-axis, right). **K**. Aggregate profile plot of Karpas-422 CXCdel H3K27me3 ChIP-Rx signal averaged across replicates (*y*-axis) centered around gene bodies of genes downregulated upon GSK343 withdrawal (*x*-axis).

**Fig. S17.**
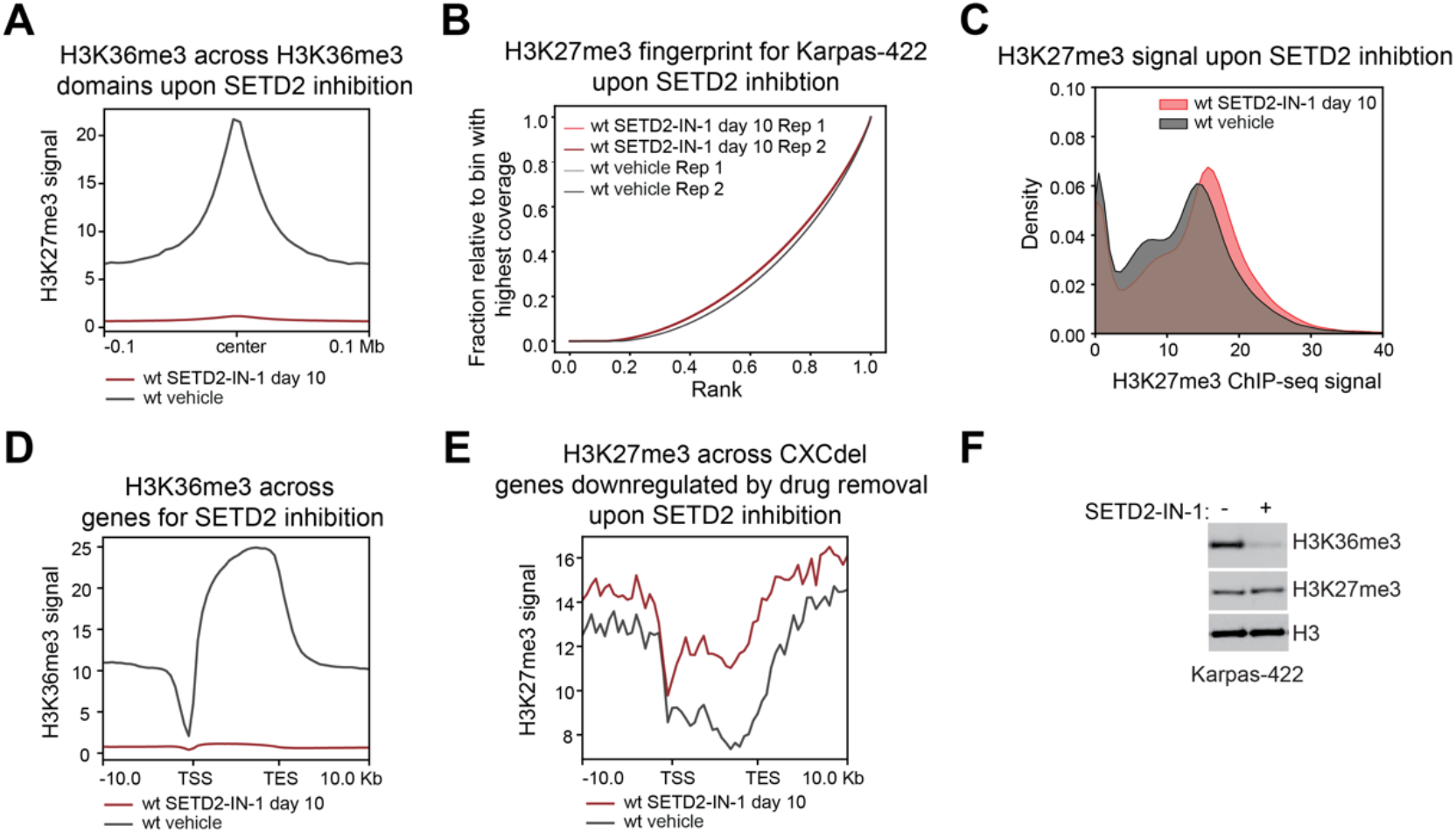
SETD2 inhibition results in modest H3K27me3 spreading on chromatin. **A**. Aggregate profile plot of wt Karpas-422 H3K36me3 ChIP-Rx signal averaged across replicates (*y*-axis) for SETD2-IN-1 (5 μM) treatment centered around H3K36me3 domains in wt Karpas-422 cells (*x*-axis). **B**. Line plot depicting fraction H3K27me3 ChIP-seq coverage relative to that of the bin with the highest coverage (*y*-axis) as a function of the region’s relative genomic rank (*x*-axis) for vehicle-treated and SETD2-IN-1-treated wt Karpas-422 cells. **C**. Density plot depicting the distribution of H3K27me3 ChIP-Rx signal averaged across replicates (*x*-axis) of SETD2-IN-1-treated wt Karpas-422 cells and vehicle-treated wt Karpas-422 cells for 50 kb genomic bins. **D**. Aggregate profile plot of Karpas-422 H3K36me3 ChIP-Rx signal averaged across replicates (*y*-axis) for SETD2-IN-1 treatment centered around gene bodies (*x*-axis). **E**. Aggregate profile plot of wt Karpas-422 H3K27me3 ChIP-Rx signal averaged across replicates (*y*-axis) for SETD2-IN-1 treatment centered around gene bodies of genes downregulated upon GSK343 withdrawal (*x*-axis). **F**. Immunoblot showing H3K27me3 and H3K36me3 levels in wt Karpas-422 under 5 μM SETD2-IN-1 treatment for 10 days.

**Fig. S18.**
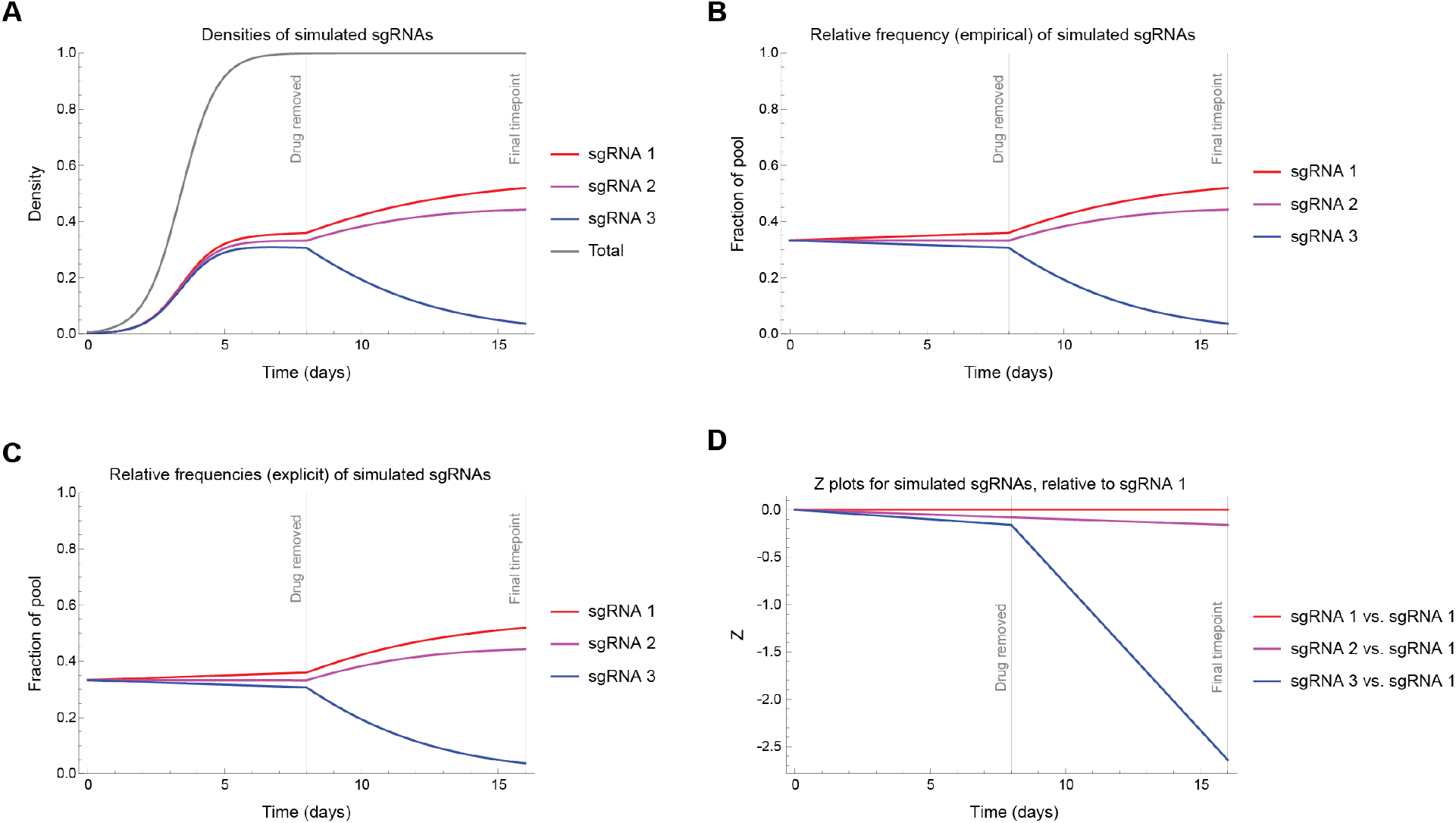
Simulated sgRNA data related to the derivation of the addiction score. **A**. Plots showing simulated densities for three hypothetical sgRNA populations. sgRNA densities begin at 1/500. The intrinsic fitnesses (uninhibited growth rates) for sgRNA populations 1–3 are 1.51, 1.50, and 1.49, respectively. When drug is removed, the fitness of sgRNA populations 1 and 2 remain unchanged, while the fitness of sgRNA 3 decreases to 1.20. The sgRNA densities over time are described by Equation 7. The total density is equal to the sum of the densities of the three sgRNAs and is described by Equation 2. **B**. Plots showing simulated sgRNA proportions according to Equation 7. Trajectories are derived by dividing the sgRNA densities depicted in **C**. 8A by the total population size. **D**. Plots showing simulated sgRNA proportions according to Equation 9. sgRNA proportions start at 0.33. The intrinsic fitness (uninhibited growth rates) for sgRNA populations 1–3 are 1.51, 1.50, and 1.49, respectively. When drug is removed, the fitness of sgRNAs 1 and 2 remain unchanged, while the fitness of sgRNA 3 decreases to 1.20. **E**. Plots showing Z-values for simulated sgRNAs relative to sgRNA population 1. The definition of Z is given in Equation 10. The slopes of these lines give may be used to compute the change in intrinsic fitness for each sgRNA when on drug vs. off drug.

**Table S1.** Proportions, Z-values, and slopes for simulated sgRNAs relative to sgRNA 1.

| **sgRNA** | **Time** | **Proportion** | **Z** | **Slope ($\sigma$)** |
| --- | --- | --- | --- | --- |
| 1 | 0 | 0.33 | 0 | — |
|  | 8 | 0.36 | 0 | 0 |
|  | 16 | 0.52 | 0 | 0 |
| 2 | 0 | 0.33 | 0.00 | — |
|  | 8 | 0.33 | -0.08 | -0.01 |
|  | 16 | 0.44 | -0.16 | -0.01 |
| 3 | 0 | 0.33 | 0.00 | — |
|  | 8 | 0.30 | -0.16 | -0.02 |
|  | 16 | 0.04 | -2.64 | -0.31 |

